# Fructose Induced KHK-C Increases ER Stress and Modulates Hepatic Transcriptome to Drive Liver Disease in Diet-Induced and Genetic Models of NAFLD

**DOI:** 10.1101/2023.01.27.525605

**Authors:** Se-Hyung Park, Robert N. Helsley, Taghreed Fadhul, Jennifer L.S. Willoughby, Leila Noetzli, Ho-Chou Tu, Marie H. Solheim, Shiho Fujisaka, Hui Pan, Jonathan M. Dreyfuss, Joanna Bons, Jacob Rose, Christina D. King, Birgit Schilling, Aldons J. Lusis, Calvin Pan, Manoj Gupta, Rohit N. Kulkarni, Kevin Fitzgerald, Philip A. Kern, Senad Divanovic, C. Ronald Kahn, Samir Softic

## Abstract

Non-alcoholic fatty liver disease (NAFLD) is a liver manifestation of metabolic syndrome, and is estimated to affect one billion individuals worldwide. An increased intake of a high-fat diet (HFD) and sugar-sweetened beverages are risk-factors for NAFLD development, but how their combined intake promotes progression to a more severe form of liver injury is unknown. Here we show that fructose metabolism via ketohexokinase (KHK) C isoform increases endoplasmic reticulum (ER) stress in a dose dependent fashion, so when fructose is coupled with a HFD intake it leads to unresolved ER stress. Conversely, a liver-specific knockdown of KHK in C57BL/6J male mice consuming fructose on a HFD is adequate to improve the NAFLD activity score and exert a profound effect on the hepatic transcriptome. Overexpression of KHK-C in cultured hepatocytes is sufficient to induce ER stress in fructose free media. Upregulation of KHK-C is also observed in genetically obesity ob/ob, db/db and lipodystrophic FIRKO male mice, whereas KHK knockdown in these mice improves metabolic function. Additionally, in over 100 inbred strains of male or female mice hepatic KHK expression correlates positively with adiposity, insulin resistance, and liver triglycerides. Similarly, in 241 human subjects and their controls, hepatic *Khk* expression is upregulated in early, but not late stages of NAFLD. In summary, we describe a novel role of KHK-C in triggering ER stress, which offers a mechanistic understanding of how the combined intake of fructose and a HFD propagates the development of metabolic complications.

## INTRODUCTION

There is a worldwide epidemic of obesity, insulin resistance, and metabolic syndrome. Non-alcoholic fatty liver disease (NAFLD) is a liver manifestation of metabolic syndrome and is estimated to affect one billion individuals worldwide (1). In the USA, NAFLD is the most common cause of chronic liver disease in children, affecting the majority of subjects with severe obesity (2) and the second most common cause of liver transplant in adults (3). In both children (4) and adults (5), NAFLD is an independent risk-factor associated with increased mortality. This disease is characterized by excessive lipid accumulation in the liver. While hepatic steatosis is a hallmark of NAFLD, lipid accumulation alone does not differentiate patients at risk of experiencing severe complications. The patients that develop liver inflammation, termed non-alcoholic steatohepatitis (NASH), are thought to be at increased risk, while those with hepatic fibrosis unequivocally have higher odds of experiencing poor outcomes. What factors mediate progression to a severe liver disease versus simple lipid deposition is the subject of active investigation.

Increased intake of dietary sugar, specifically fructose, has been associated with development of severe NAFLD. Adult subjects with NAFLD consume twice as many calories from sugar-sweetened beverages as do the subjects with non-steatotic livers (6). Moreover, increased fructose consumption is positively correlated with the severity of liver fibrosis in adults with NAFLD (7). In children, fructose consumption is an independent risk-factor for NASH (8). This is concerning as adolescents consume more sugar than any other age group. We showed that the first enzyme of fructose metabolism, ketohexokinase (KHK), is increased in liver biopsies from adolescents with NASH, as compared to obese children with normal livers (9).

The only known function of KHK is to metabolize fructose to fructose-1 phosphate. There are two main KHK isoforms, KHK-A and KHK-C, produced by alternative splicing of a common gene (10). KHK-A is expressed ubiquitously, while KHK-C is found predominantly in the liver, kidney and intestine, and is the primary isoform responsible for fructose metabolism (11). In agreement with its only known function, KHK-C is upregulated by fructose intake. More recently, it has been found that KHK-C is also induced by hypoxia (12), as fructose produced endogenously from glucose can serve as an alternative energy source in an environment that does not support normal glucose metabolism (13).

The detrimental metabolic effects of fructose are thought to be secondary to its high lipogenic potential (14). However, fructose-induced lipogenesis is the highest on a chow diet and decreases on a HFD (9), since diets high in fat provide ample supply of lipids. In spite of strongly inducing lipogenesis on chow diet, metabolic effects of dietary fructose are best observed in the setting of combined intake with a HFD (15). HFD supports development of metabolic complications, in part, via propagation of ER stress. This is an adaptive cellular response aimed at maintaining energy homeostasis and is mediated through three pathways, including activating transcription factor 6 (ATF6), inositol-requiring transmembrane kinase – endoribonuclease 1 activating x-box binding protein 1 (IRE1a-XBP1) and protein kinase RNA-like endoplasmic reticulum kinase (PERK) (16). On the other hand, unresolved ER stress triggers a maladaptive response that supports development of metabolic complications, including a severe form of NAFLD (17).

In this study, we have investigated the combined intake of fructose-sweetened water and a HFD on the development of severe fatty liver disease. We find that fructose-induced KHK-C, via its previously unrecognized role, induces ER stress. When combined with a HFD intake this results in ineffective signal transduction via the PERK arm of ER stress to accelerate the development of glucose dysregulation, liver inflammation and fibrosis. A liver-specific knockdown of KHK is sufficient to reverse metabolic dysregulation in mice consuming fructose on a HFD, but also in mice with genetic causes of obesity or lipodystrophy, on a fructose free diet. The broad effects of KHK knockdown are likely mediated via its reduction of ER stress.

## RESULTS

### Consumption of fructose on a HFD leads to unresolved ER stress

A cohort of 10-week-old mice were fed chow or HFD and given *ad libitum* access to regular water or water containing 30%(w/v) fructose for 16 weeks. Mice supplemented with fructose (Fruct) on a chow diet gained more weight (36.0±1.1g) than the mice on a chow diet (Chow) drinking regular water (32.7±1.3g) (Fig S1A). HFD-fed mice drinking regular water (HFD) gained more weight (46.6±1.1g) than both chow-fed controls. However, supplementation with fructose on a HFD (HFD+F) did not result in additional weight gain (46.9±0.9g). In agreement with weight gain, subcutaneous (SQ), visceral adipose tissue (VAT) and brown adipose tissue (BAT) were generally increased in Fruct compared to the Chow group (Fig S1B). HFD-fed mice had even higher increase in adipose tissue mass, while HFD+F-fed mice did not have further increase in adiposity, compared to the HFD group. Similarly, liver weight was increased in mice on HFD (1.8±0.2g) compared to the Chow (1.2±0.1g), but it was not significantly increased by addition of fructose to a HFD (2.1±0.2g). The total caloric intake was increased in mice on every diet compared to the Chow alone, but there was no difference among the hypercaloric diets (Fig S1C). The mice consuming fructose-sweetened water had corresponding decrease in solid food intake on both chow and HFD, since 43% to 24% of their calories came from drinking fructose.

Fructose intake on a HFD accelerates liver injury in murine models of fatty liver disease (18; 19). Serum alanine aminotransferase (ALT), a marker of liver injury, was comparable in mice in Chow and Fruct groups, but was significantly higher (p<0.01) in mice on a HFD and addition of fructose to a HFD further elevated ALT (p<0.0001) (Fig 1A). Serum triglycerides were not increased by the diets (Fig S1D), while LDL cholesterol was higher in the Fruct group compared to Chow and in both HFD and HFD+F groups. VLDL and HDL cholesterol were not different (Fig S1E). Marked steatosis developed in mice on a HFD and it was exacerbated with addition of fructose, leading to more severe micro and macrovesicular steatosis (Fig 1B). Liver triglycerides were not increased in mice on Fruct, as compared to the Chow group (Fig 1C). HFD-fed mice had higher levels, while addition of fructose did not further raise hepatic triglycerides. This data agrees with our previous findings indicating that lipid composition, rather than total triglyceride accumulation, correlates with progression to a more severe form of fatty liver disease (9).

**Figure 1.**
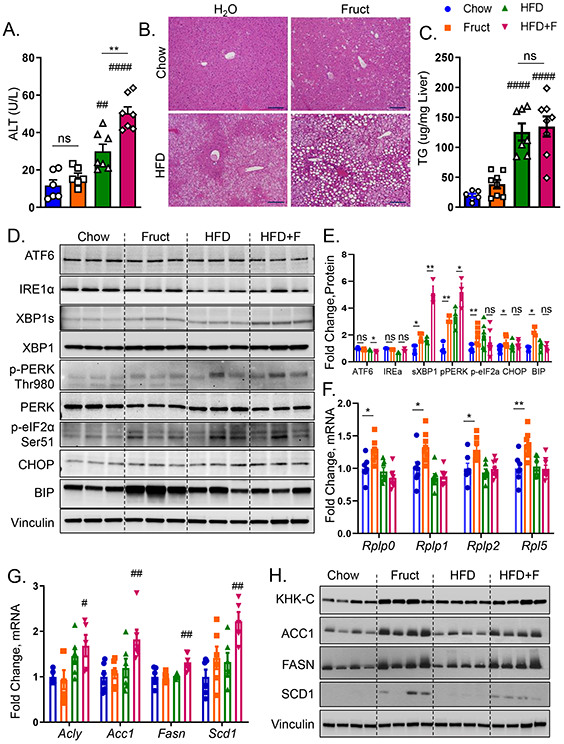
Intake of fructose and HFD leads to unresolved ER stress. A) Serum alanine aminotransferase (ALT) at sacrifice in male, C57BL/6J, mice drinking regular water (Chow) or 30% fructose-sweetened (Fruct) water on Chow diet or regular (HFD) or fructose water (HFD+F) on a HFD for 16 weeks. n=6-8 mice per group. B) Representative liver H&E histology, scale bar = 100 μm C) hepatic triglycerides from these mice. D) Western blot of (ATF6, IRE1α, XBP1, PERK, eIF2α, BIP, CHOP) and E) Image J quantification of their protein levels. n=3-6 per group. F) mRNA expression of ribosomal subunits (*Rplp0*, *Rplp1*, *Rplp2*, *Rpl5*), as well as G) mRNA and H) western blot of lipogenic enzymes (ACLY, ACC1, FASN, SCD1) from liver homogenates of these mice. Sample number for WB is n=4 and for mRNA n=6 mice per group. Statistically significant results are marked with # using one-way ANOVA analysis, with Dunnett’s multiple comparisons test to the Chow group, # p<0.05; ## p<0.01; #### p<0.0001, while * represents a post-hoc analysis between the groups under the line, * p<0.05; ** p<0.01. All data are presented as mean ± S.E.M.

Three distinct pathways – ATF6, IREα-XBP1 and PERK mediate endoplasmic reticulum (ER) stress. ER stress has been proposed to drive liver injury and progression to NASH (17). Western blot analysis of liver homogenates revealed that the ATF6 arm of ER stress was not increased by fructose containing diets (Fig 1D). Similarly, IRE1α was not affected, but downstream spliced XBP1 was increased 2-fold in Fruct group compared to the Chow (Fig 1D, 1E). XBP1s was also increased to a similar level in mice on a HFD, and it was further elevated ~5-fold in HFD+F group. It has been reported that PERK pathway plays an important role to enhance signaling by the IRE1α to increase Xbp1s (20). Phosphorylated PERK at Thr980 was increased ~3-fold in both Fruct and HFD groups, and it was stepwise 5.2-fold higher in HFD+F group (Fig 1D, 1E). We also assessed pPERK by IHC and it was elevated in the Fruct and HFD groups compared to the Chow, but was the highest in the HFD+F group (Fig S1F). Activated PERK propagates ER stress by phosphorylating eukaryotic translation initiation factor 2a (eIF2α). Indeed, eIF2α was phosphorylated at Ser51 in both Fruct and HFD groups, but addition of fructose to a HFD failed to further propagate eIF2α phosphorylation. Similarly, apoptotic marker, C/EBP homologous protein (CHOP) was increased in Fruct, but was not further significantly elevated in HFD+F group. ER chaperone, binding immunoglobulin protein (BIP) was elevated in Fruct group, but it actually decreased in HFD+F group in spite of upstream elevation in pPERK arm of ER stress consistent with unresolved ER stress in this group (Figs 1D, 1E). Phosphorylated eIF2α upregulates transcription of enzymes participating in protein folding, while at the same time it attenuates protein translation through mRNA degradation (21). Indeed, mRNA levels of protein folding chaperones *Hspa5, Edem1* and Hsp90b1were increased in Fruct group compared to the Chow, but were not further induced in HFD+F group compared to the HFD (Fig S1G). Similarly, mRNA expression of genes coding ribosomal 60S subunit, such as *Rplp0, Rplp1, Rplp2* and *Rpl5* that support protein translation, was increased in Fruct, but not in HFD+F group (Fig 1F). Conversely, the expression of genes regulating *de novo* lipogenesis (DNL) including *Acly, Acc1, Fasn* and *Scd1* were not increased in Fruct group, but their expression was increased in HFD+F (Fig 1G). Interestingly, protein levels of these same DNL enzymes were increased with fructose supplementation of both Chow and HFD (Figs 1H, S1H) suggestive of post-translational protein stabilization. The discrepancy between mRNA and protein levels may indicate that indeed increased phosphorylation of eIF2α in Fruct group leads to selective mRNA degradation, a known function of phosphorylated eIF2α (21). Ineffective phosphorylation of eIF2α did not induce mRNA decay of DNL genes in HFD+F group. In summary, both fructose and HFD induced XBP1s and PERK arms of ER stress, but when these nutrients were consumed together, this led to ineffective signaling via the PERK pathway leading to unresolved ER stress.

### The combined intake of fructose and HFD worsens hepatic insulin resistance, inflammation and fibrosis

ER stress can lead to metabolic dysfunction, so we measured glucose tolerance in our mice after 8 weeks on the diets. Fruct group had normal glucose tolerance, similar to the Chow (Figs S2A, S2B). HFD-fed mice did not develop glucose intolerance at this early time point. However, the combined intake of fructose and a HFD impaired glucose tolerance. Whole body insulin sensitivity was also not impaired in Fruct, as compared to Chow group, but both HFD-fed groups developed insulin resistance (Figs S2C, S2D). However, when insulin levels were measured over time, insulin increased sooner and rose higher in HFD+F group, compared to the HFD alone (Fig S2E), even though at the end of the experiment serum insulin was not statistically different between the groups.

Glucose tolerance is a product of endogenous glucose production, predominantly made by the liver and glucose utilization by almost every tissue. Insulin sensitivity is largely driven by insulin-stimulated glucose uptake in the muscle and adipose tissue. Since these tissues do not metabolize fructose, the difference between GTT and ITT could be explained by hepatic insulin resistance. Next, we assessed hepatic insulin signaling following insulin or saline injection via inferior vena cava 10 minutes before sacrifice. Compared to saline treated mice, AKT S473 phosphorylation was robustly increased in Chow group injected with insulin, documenting normal hepatic insulin signaling (Fig S2F). Fruct and HFD-fed groups had slightly lower p-AKT, whereas HFD+F group showed profound decrease in p-AKT. ERK phosphorylation increased in all four insulin-treated groups, documenting technically adequate insulin injection. The magnitude of ERK phosphorylation was lower in HFD+F group, consistent with decreased AKT phosphorylation. In line with AKT and ERK results, pan-tyrosine phosphorylation of insulin signaling molecules was decreased in Fruct and HFD groups and was the lowest in HFD+F group documenting severe hepatic insulin resistance. To further probe the mechanism of insulin resistance we quantified protein levels of several important mediators of insulin signaling. Insulin receptor (IR) was reduced in HFD and HFD+F groups compared to the Chow (Figs S2G, S2H). This is consistent with hyperinsulinemia in these mice, since insulin is known to downregulate its receptor (22). Interestingly, insulin receptor substrate 1 (IRS-1) tended to be reduced in Fruct group, but it was significantly reduced in HFD+F group, compared to the Chow. A regulatory subunit of phosphatidylinositol 3-kinase, p85α, was not affected by the diets.

In addition to impaired insulin signaling, ER stress may propagate NAFLD progression by supporting inflammation via c-Jun NH_2_-terminal kinase (JNK) pathway. Indeed, fructose supplementation of either Chow or HFD significantly amplified JNK Thr183/Tyr185 phosphorylation (Fig S3A). In agreement with activation of JNK pathway, hepatic mRNA expression of macrophage markers documented increased *Cd11c, F4/80*, and *Mip* in HFD+F group (Fig S3B). Moreover, mRNA expression of genes involved in fibrogenesis, such as *Tgf*β, α*Sma*, and *Col1a* were significantly increased in HFD+F fed mice, compared to the Chow (Fig S3C). Sirius red stain confirmed increased collagen deposition in HFD+F-fed mice (Fig S3D). These data are in agreement with published studies showing that addition of fructose to a HFD increases liver inflammation and fibrosis (23; 24).

### Knockdown of T-KHK in mice on HFD+F-diet improves metabolic dysfunction by decreasing de novo lipogenesis and ER stress

Utilizing liver-specific siRNA made by Alnylam Pharmaceuticals, we knocked down (KD) total KHK (T-KHK) in the livers of male, C57BL/6J mice beginning after 12 weeks of HFD+F feeding. Following 8 weeks of bimonthly siRNA injections, mice were sacrificed after 20 weeks on the diets. Expression of *Khk-c* and *Khk-a* isoforms was elevated in HFD+F-fed mice compared to the Chow (Fig 2A). KD of T-KHK decreased both *Khk-c* and *-a* mRNA better than 95 percent. Whereas both KHK isoforms were decreased, *Khk-c* mRNA is about 30 times more abundant in the liver then *Khk-a* (Fig S4A). Similarly, mRNA expression of enzymes catalyzing the second and third steps of fructose metabolism (Fig S4B) were increased in HFD+F, compared to the Chow group (Fig 2B). T-KHK KD had no effect on the expression of these fructolytic genes, confirming that we specifically knocked down the first step of fructose metabolism. Hepatic fructose metabolism can be upregulated by endogenous fructose produced, via the polyol pathway, from glucose as a substrate (11). Aldose reductase (*Akr1b1*), which converts glucose to sorbitol, and sorbitol dehydrogenase (*Sord*), which converts sorbitol to fructose were unaffected by fructose supplementation or T-KHK KD (Fig 2C).

**Figure 2.**
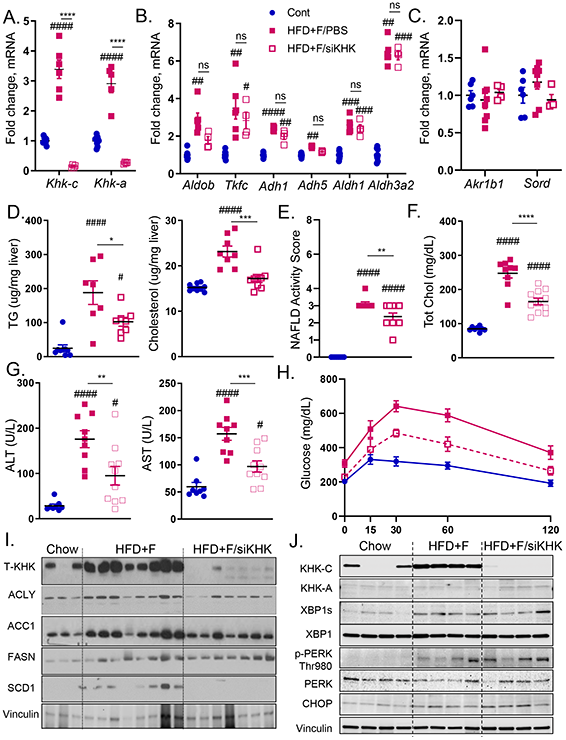
Ketohexokinase knockdown improves metabolic dysfunction in HFD+F-fed mice. A) mRNA expression of genes regulating first, as well as B) second and third steps of fructose metabolism and genes responsible for C) endogenous fructose production in the livers of male, C57BL/6J mice fed chow diet on regular water (Chow), or HFD supplemented with 30% fructose and injected with PBS (HFD+F/PBS) or injected with KHK siRNA (HFD+F/siKHK) for the last 8 out of 20 weeks on the diets. Chow group for every gene was normalized to 1. Hepatic D) triglycerides (TG) and total cholesterol levels in the livers of these mice. E) NAFLD activity score based on the interpretation of liver histology by an independent laboratory. Serum F) total cholesterol, as well as G) alanine aminotransferase (ALT) and aspartate aminotransferase (AST) obtained at sacrifice, n=8-11 mice per group. H) Glucose tolerance test in these mice performed after 18 weeks on the diet. Western blot of liver KHK and proteins mediating I) *de novo* lipogenesis (ACLY, ACC1, FASN, SCD1) and J) ER stress (XBP1, PERK, CHOP) pathways. Sample number for WB is n=3-8 and for mRNA n=6 mice. Statistically significant results are marked with # using one-way ANOVA analysis, with Dunnett’s multiple comparisons test to the Chow group, # p<0.05; ## p<0.01; ### p<0.001; #### p<0.0001, while * represents a post-hoc analysis between the groups under the line, *p <0.05; ** p<0.01, *** p<0.001, **** p<0.0001. All data are presented as mean ± S.E.M.

Body weight in HFD+F group (50.6±1.3g) was significantly higher than in the Chow group (32.1±1.3g), and liver-specific T-KHK KD decreased weight gain (46.3±1.4g) (Fig S4C). Consistent with weight, liver mass was higher in HFD+F group (2.7±0.3g) than in Chow (1.5±0.1g), and T-KHK KD lowered liver weight by 20% (2.1±0.2g) (Fig S4D). Similarly, hepatic triglycerides and cholesterol were reduced following T-KHK KD (Fig 2D). Liver histology was graded by Experimental Pathology Laboratories, Inc., an independent company hired to objectively grade liver histology. NAFLD activity score (NAS) consists of steatosis grade (0-3), inflammation (0-3), and balloon degeneration (0-2) (25). NAS was elevated in HFD+F group and it meaningfully (p<0.01) improved following T-KHK KD (Fig 2E). This improvement was primarily driven by a decline in steatosis grade (Fig S4E), whereas inflammation was unchanged (Fig S4F) and none of the mice had hepatocyte ballooning. Next, we quantified serum markers of metabolic dysfunction. Total cholesterol (Fig 2F), as well as HDL and LDL cholesterol (Fig S4G), were elevated in mice fed a HFD+F diet and they markedly improved (p<0.001) following KD of T-KHK. Serum markers of liver injury, ALT and AST, were also elevated in HFD+F group and they significantly declined in HFD+F/siKHK group (Fig 2G). Other markers of liver function, alkaline phosphatase and total bilirubin, were unaffected by the diets or T-KHK KD (Fig S4H). Basal glucose levels were increased in HFD+F mice and they normalized in HFD+F/siKHK group (Fig S4I). Similarly, glucose tolerance was impaired in HFD+F group, compared to the Chow, whereas T-KHK KD markedly (p<0.001) improved glucose tolerance in HFD+F/siKHK mice (Figs 2H, S4J).

To interrogate the mechanism underlying improved metabolic dysfunction following T-KHK KD, we first assessed the *de novo* lipogenesis pathway. Enzymes mediating DNL, such ACLY, ACC1, FASN and SCD1 (14), were increased 5-10 fold in HFD+F-fed mice compared to the Chow (Figs 2I, S4K). These proteins normalized following a T-KHK KD to the level observed in Chow group. An increase in DNL proteins correlated with higher T-KHK, which is mainly driven by an increase in KHK-C, rather than KHK-A isoform (Figs 2I, 2J). Next, we quantified proteins involved in mediating the two arms of ER stress that were altered by dietary fructose. XBP1s was increased in HFD+F-fed mice compared to the Chow, but KD of T-KHK did not affect XBP1s (Figs 2J, S4L). On the other hand, PERK arm of ER stress was increased in mice on HFD+F diet, and T-KHK siRNA treatment further increased PERK phosphorylation. Similarly, CHOP, a downstream target of activated PERK, was increased in mice on a HFD+F diet, and CHOP significantly increased following a T-KHK KD. These data suggest that both XBP1s and PERK pathways of ER stress are affected by dietary fructose, but only PERK-CHOP pathway was increased by a KD of T-KHK. In summary, liver-specific KD of T-KHK in mice consuming a HFD+F diet was associated with improved hepatic and whole body metabolic dysfunction, which was accompanied by decreased DNL and improved PERK-CHOP arm of ER stress.

### Knockdown of T-KHK profoundly alters transcriptional activity in the liver

To comprehensively investigate metabolic pathways affected by fructose supplementation of a HFD and by T-KHK KD we performed RNAseq analysis. Principal component analysis showed that liver transcriptome of a HFD+F group was substantially different from that of the Chow group (Fig 3A). Interestingly, KD of T-KHK segregated from the other groups in the PCA plot, illustrating its transformation of the liver transcriptome (Fig 3A). This is an interesting findings since the only known function of KHK is to phosphorylate fructose. Next, we identified the most significantly altered pathways from this analysis. Consistent with a well-known function of ER stress to halt transcriptional activity, the most significantly upregulated pathway was transcriptional regulation by E2F transcription factor 6 (E2F-6) (Fig 3B). The protein encoded by E2F-6 gene contains a modular suppression domain that functions to inhibit transcription (26). Additionally, ChIP-seq from the ENCODE Transcription Factor Targets dataset identified eIF2α as a target gene of E2F-6. Other significantly altered pathways included nuclear estrogen signaling (Estrogen Dependent Nuclear Events pathway and Extra Nuclear Estrogen Signaling pathway), as well as nucleotide recycling (Nucleotide Salvage pathway, Metabolism of Nucleotides pathway, Nucleobase Catabolism pathway and Pyrimidine Catabolism pathway). Nucleotide salvage pathways recycle nucleotides formed during degradation of RNA and DNA. Again, this is likely a consequence of increased ER stress as it promotes mRNA degradation to decrease transcription. In congregate, seven out of the ten most significantly altered pathways are involved in regulation of transcription. Alternatively, E2F6 and nucleotide metabolism pathways can be involved in cell cycle regulation, which may be altered by KHK knockdown.

**Figure 3.**
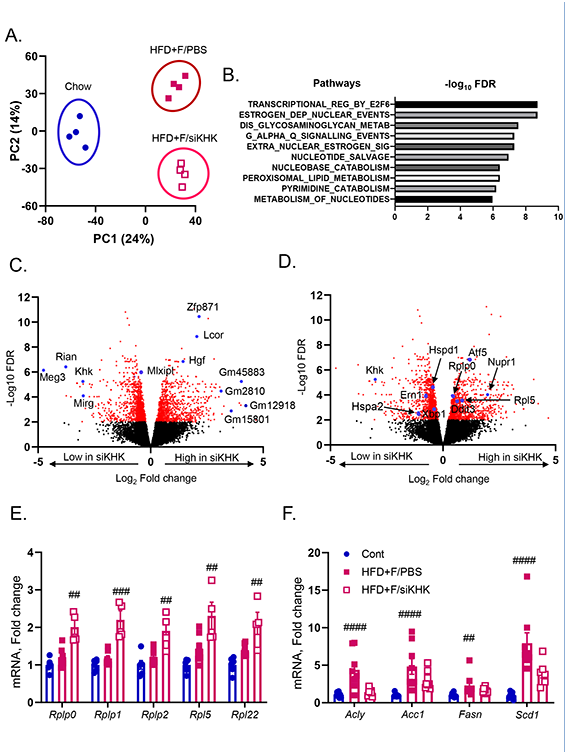
Knockdown of T-KHK in mice on HFD+F diet profoundly alters hepatic transcriptome. A) Principal component analysis (PCA) of RNAseq data from the livers of mice presented in figure two. B) The most significantly altered pathways from this analysis. Volcano plot analysis in HFD+F-fed mice showing upregulated and downregulated genes with KHK knockdown (siKHK) compared to HFD+F/PBS control. Some of the most altered genes are labeled in C) and in D) are labeled the genes involved in ER stress. mRNA expression of genes regulating E) ribosomal subunits (*Rplp0*, *Rplp1*, *Rplp2*, *Rpl5*) and de novo lipogenesis (*Acly, Acc1, Fasn, Scd1*).

Volcano plot analysis comparing HFD+F versus HFD+F/siKHK groups showed that indeed one of the most significantly downregulated genes with T-KHK KD is KHK, but also maternally expressed gene 3 (*Meg3*), RNA imprinted and accumulated in nucleus (*Rian*), and miRNA containing gene (*Mirg*) (Fig 3C). *Meg3*, *Rian* and *Mirg* are closely linked maternally expressed non-coding RNAs that function as an integral part of genome-wide regulatory network and have been implicated in ER stress (27; 28). On the other hand, some of the most significantly upregulated genes with T-KHK KD are RNA binding protein, zinc finger protein 871 (*Zfp871*), ligand dependent nuclear receptor corepressor (*Lcor*) and hepatocyte growth factor (*Hgf*), as well as many long non-coding RNAs such as *Gm45883*, *Gm2810*, *Gm12918* and *Gm15801* (Fig 3C). Narrowing down on the ER stress response, downregulated genes with T-KHK KD include *Xbp1* and *Ern1* (aka *Ire1a)*, as well as heat shock proteins *Hspd1* and *Hspa2* (Fig 3D). Upregulated genes with T-KHK KD include DNA damage inducible transcript 3 (*Ddit3*), also known as *Chop*, its target, activating transcription factor 5 (*Atf5*) and nuclear protein 1 (*Nupr1*) required for eIF2α mediated resolution of ER stress, as well as ribosomal subunits *Rplp0* and *Rpl5*. In figure 1E we showed that upregulation of these ribosomal subunits was impaired in HFD+F-fed mice, so we verified this part of RNAseq analysis via qPCR. Consistent with our previous results, mRNA expression of *Rplp0, Rplp1, Rplp2, Rpl5* and *Rpl22* was not increased in HFD+F-fed mice compared to the Chow, but they were increased following T-KHK KD (Fig 3E). We also quantified DNL pathway by qPCR and it was increased in HFD+F but not in mice treated with KHK siRNA (Fig 3F).

Heatmap analysis again showed that the most significantly upregulated genes in HFD+F-fed mice were non-coding RNAs *Mirg, Rian* and *Meg3*, whereas the most significantly increased genes in HFD+F/siKHK group were genes regulating transcription (*Zfp871, Lcor*) and long non-coding RNAs (Fig S5A). Lipogenesis pathway was also affected by T-KHK KD, so that the genes regulating fatty acid synthesis pathway such as *Mlxpl* (aka *Chrebp*), *Pklr, Scd1* and *Acaca* (aka *Acc1*) were downregulated, whereas genes regulating triglyceride assembly such as *Agpat1,2*, *Gpat3* and *CideC* were upregulated with T-KHK KD (Fig S5B). In summary, KD of T-KHK had a large and unanticipated effect on global hepatic transcriptional activity, and more specifically on DNL and ER stress pathways.

### Overexpression of KHK-C is sufficient to upregulate ER stress in vitro

In light of the profound effect of T-KHK KD on ER stress and thus hepatic transcriptome, we overexpressed (OE) GFP tagged (27 kDa) mouse KHK-C in human HepG2 cells cultured in standard (DMEM), fructose-free media. We (15; 29) and others (30) have previously published that HepG2 cells do not express KHK-C. KHK-C in HepG2 cells was overexpressed to the level similar to HFD+F-fed mice (Fig 4A). Remarkably, OE of KHK-C was sufficient to increase *Ddit3 and* ER chaperones *Hspa5, Edem1* and *Hsp90b1* (Fig 4B). Furthermore, ribosomal subunits *Rplp1, Rplp2, Rpl5* and *Rpl22* were also increased simply by KHK-C OE (Fig 4C).

**Figure 4.**
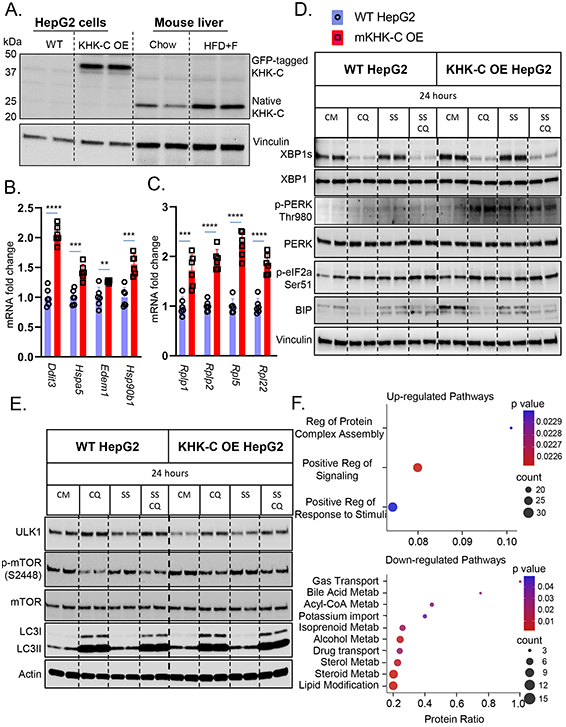
Overexpression of KHK-C Promotes ER Stress in vitro. Wild type (WT) HepG2 hepatocytes were engineered to overexpress (OE) mouse KHK-C (mKHK-C OE) tagged with GFP. A) Western blot for KHK-C in WT and KHK-C OE cells as compared to liver homogenates from Chow and HFD+F-fed mice from Fig1. mRNA expresses as assessed by qPCR of B) ER-stress (*Ddit3*, *Hspa5, Edem1*, *Hsp90b1*), C) ribosomal (*Rplp1*, *Rplp2*, *Rpl5*, *Rql22*) genes were measured in WT HepG2 and mKHK-C OE cells, n=6 wells per group. D) WT HepG2 and mKHK-C OE hepatocytes were treated with complete media (CM), chloroquine (CQ), serum-starvation (SS), and a combination of serum starvation and chloroquine for 24 hours. Western blot was used to analyze D) ER-stress and E) autophagy pathways. mKHK-C was also OE in AML-12 mouse hepatocytes to minimize cell specific effects. Proteomic analysis using data-independent acquisitions on an Orbitrap Eclipse Tribrid platform was performed and most significantly F) upregulated and downregulated pathways were determined using ConsensusPathDB functional annotation pathway analysis. Statistics were completed by an unpaired Student’s t-test for qPCR data. \*\**P*≤0.01; \*\*\**P*≤0.001; \*\*\*\**P*≤0.0001. All data are presented as mean ± S.E.M.

Next, we quantified ER stress pathway in WT and KHK-C OE HepG2 cells by western blot. The cells were treated with or without chloroquine (CQ), to prevent autophagosome-lysosome fusion, and were serum starved (SS), to further induce ER stress (Fig 4D). In WT cells, CQ treatment profoundly decreased XBP1s, confirming a link between autophagy and ER stress. Serum starvation had no effect on XBP1s in WT cells. Remarkably, OE of KHK-C increased XBP1s protein, and again, CQ decreased it, while SS had no effect. PERK phosphorylation at Thr980 was not profoundly affected by CQ or SS in WT cells. OE of KHK-C increased p-PERK compared to WT cells. CQ treatment further increased it, whereas SS, adding a second form of ER stress, increased PERK phosphorylation. Downstream eIF2α phosphorylation was increased by CQ or SS exposure in WT cells. KHK-C OE showed increased p-eIF2α compared to WT and it was further increased by CQ treatment. SS also increased p-eIF2α, but to a lesser extent than CQ exposure. We could not detect CHOP in this cancer cell line, so we quantified BIP protein. CQ treatment decreased, while SS increased BIP levels in WT cells. KHK-C OE cells had markedly elevated BIP in line with increased ER stress in these cells. Remarkably, adding a second form of ER stress in terms of SS actually decreased BIP, similar to when fructose was added to a HFD (Fig 1D).

KHK-C OE HepG2 cells were cultured in fructose free media, but to ensure that KHK-C induces ER stress independent of fructose metabolism we also OE mutant kinase dead KHK-C (KHK-C MD). Mutant kinase dead KHK-C was generated by point mutation (G527R) in ATP binding domain so that the enzyme can bind, but cannot phosphorylate fructose. Compared to GFP OE cells, OE of either WT or MD KHK-C increased PERK phosphorylation indicative of elevated ER stress (Fig S6A). Additionally, KHK-C WT enzyme was OE in chow-fed C57BL/6J mice for one week using adeno associated virus (AAV), serotype 8. Compared of GFP, OE of KHK-C in vitro induced PERK and eIF2α phosphorylation and increased CHOP protein levels (Fig S6B).

ER stress regulates autophagy (31), an intracellular degradation pathway for recycling improperly folded proteins (32). Autophagy initiation is mediated by the unc-51 like autophagy activating kinase 1 (ULK1) and is inhibited by mammalian target of rapamycin (mTOR). ULK1 was decreased, while phosphorylation of mTOR at S2448 was increased in KHK-C OE cells (Fig 4E), consistent with decreased autophagy. LC3 mediates the final step of autophagosome fusion with lysosome and consists of native, LC3-I, and lipidated, active form LC3-II. CQ blocks autophagy flux enabling steady state inference. In CQ treated WT cells, the ratio of LC3 II/I was increased with SS indicative of increased autophagy. On the other hand, LC3 II/I ratio was decreased in CQ and CQ+SS treated KHK-C OE cells consistent with decreased autophagy flux. This is primarily evident by increased inactive LC3-I in KHK-C OE cells. Taken together, these data indicate that increased ER stress in KHK-C OE hepatocytes leads to decreased autophagy flux, a well-known role of unfolded protein response.

Next, we asked if decreased autophagy flux is accompanied with increased ER-associated degradation of unique proteins mediating insulin signaling. WT and KHK-C OE HepG2 cells were treated with Cycloheximide (Chx) to prevent protein synthesis and treated with fructose (F) or CM for 4 or 8 hours. IRS-1 was rapidly degraded in WT cells treated with fructose and its degradation was enhanced in KHK-C OE cells (Fig S6C), in agreement with decreased IRS-1 with fructose treatment *in vivo* (Fig S2G). AKT and ERK proteins were minimally decreased after 8 hours of Chx treatment, likely due to their longer half-life, but this decrease was stronger in F-treated KHK-C OE cells. On the other hand, p85α node of insulin signaling was unaffected by Chx treatment similar to no change in p85α protein *in vivo* (Fig S2G).

We also asked if PERK arm of ER stress can be degraded by KHK-C OE in HepG2 cells. Again PERK, p-eIF2a and BIP were higher in KHK-C OE cells compared to WT (Fig S6D). Total PERK levels were lower in F- compared to CM-treated cells after 4h of Chx treatment in KHK-C OE, but not WT cells (Fig S6D). However, p-eIF2α and BIP were not different in F-, compared to CM-treated cells. In summary, KHK-C OE leads to increased ER stress and ineffective PERK phosphorylation when exposed to additional cellular stress. ER stress may cause ApoB degradation that feeds into hepatic steatosis (33). Indeed, ApoB was decreased in HFD+F group (Fig S6E), consistent with severe hepatic steatosis in these mice.

To comprehensively evaluate OE of KHK-C in a different cell type, we performed proteomic analysis in AML-12 mouse hepatocytes cultured in fructose-free media using data-independent acquisitions (34; 35) on an Orbitrap Eclipse Tribrid platform. We identified and quantified 4,252 unique proteins. Only the proteins with two or more unique peptides were quantified in order to increase specificity. Four hundred and four proteins were significantly altered comparing KHK-C OE vs WT cells (q-value <0.01 and log2(fold change)>0.58). Of these 58.9% were downregulated and 41.1% were upregulated. ConsensusPathDB functional annotation analysis revealed only three upregulated pathways, and the most significant one was ER dependent Regulation of Protein Complex Assembly (Fig 4F). The top ten downregulated pathways included processes involved in acyl-CoA, cholesterol, and lipid metabolism, consistent with metabolic dysfunction observed in the setting of ER stress (Fig 4F).

### Knockdown of T-KHK in ob/ob mice improves metabolic dysfunction and ER stress, but not DNL

In the figures above, we have described a novel function of KHK-C in ER stress. Next, we tested whether T-KHK KD has an effect in mice with genetically induced obesity and ER stress, independent of dietary fructose intake. Two-month-old, chow-fed, leptin-deficient (ob/ob) mice were treated with T-KHK or luciferase control siRNA for four weeks. Weight gain in ob/ob mice was significantly greater (12.5±0.8g) than in heterozygote (ob/+ Het) control mice (4.6±0.8g) (Fig 5A). T-KHK KD significantly (p<0.01) reduced weight gain in ob/ob-siKHK (9.1±0.9g), compared to the ob/ob-siCont group. Reduced weight gain is reflected, in part, by 20% reduction in liver weight in ob/ob-siKHK group (4.0±0.1g), compared to ob/ob-siCont (3.4±0.2g) (Fig 5B). This is accompanied by a trend (p=0.08) toward reduced hepatic triglycerides, whereas hepatic cholesterol was significantly (p<0.01) lower in ob/ob-siKHK group (Fig 5C). Furthermore, perigonadal VAT (Fig S7A) was lower, while kidney weight (Fig S7B) was unchanged between ob/ob-siKHK and ob/ob-siCont groups.

**Figure 5.**
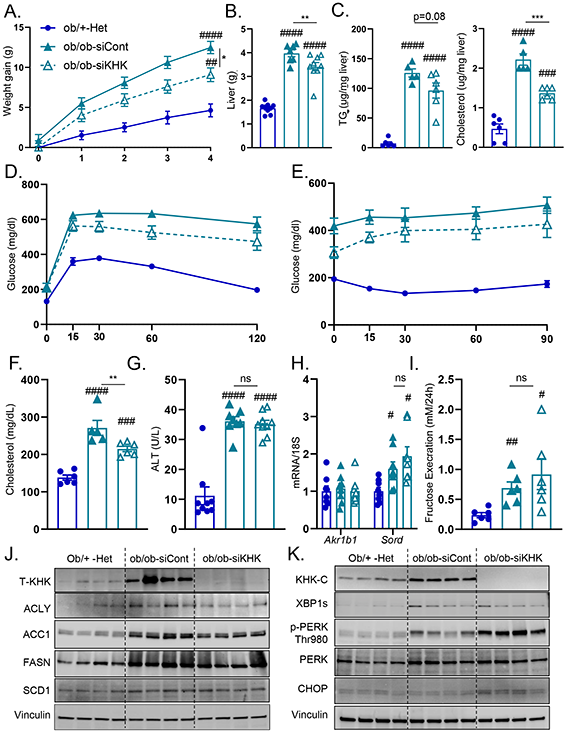
ER stress, but not lipogenesis, improves following T-KHK knockdown in Ob/Ob mice. Weight gain A) in 2-month-old, male, heterezygote (Ob/+-Het), as well as in homozygous, leptin-deficient mice (Ob/Ob-siCont) treated with luciferase control, or KHK siRNA (Ob/Ob-siKHK) for four weeks, n=6-9 mice per group. B) Liver weight, C) hepatic triglycerides (TG) and cholesterol levels in these mice. D) Glucose tolerance test in these mice performed after 2 weeks following siRNA injection. E) Insulin tolerance test in these mice performed after 3 weeks following siRNA injection. Serum F) total cholesterol, as well as G) alanine aminotransferase (ALT) obtained at sacrifice, n=6-9 mice per group. Hepatic mRNA expression of H) genes regulating endogenous fructose production (*Akr1b1, Sord*). n=6 mice per group. I) Urinary fructose determined following 24h urine collection. Western blot of liver lysates for KHK and proteins mediating J) *de novo* lipogenesis (ACLY, ACC1, FASN, SCD1) and K) ER stress (KHK-C, XBP1, PERK, CHOP) pathways. Sample number for WB is n=4. Statistically significant results are marked with # using one-way ANOVA analysis, with Dunnett’s multiple comparisons test to the Chow group, # p<0.05; ## p<0.01; ### p<0.001; #### p<0.0001, while * represents a post-hoc analysis between the groups under the line, *p <0.05; ** p<0.01, *** p<0.001, **** p<0.0001. All data are presented as mean ± S.E.M.

Next, we assessed metabolic dysfunction in these mice. Glucose tolerance was impaired in ob/ob mice compared to the heterozygote controls (Figs 5D, S7C), and T-KHK KD significantly (p<0.01) improved glucose tolerance. Similarly, both ob/ob-siKHK and ob/ob-siCont mice had impaired insulin sensitivity, and T-KHK KD tended to improve insulin sensitivity (Figs 5E, S7D). Serum cholesterol was elevated in ob/ob mice and it significantly (p<0.01) improved with T-KHK KD (Fig 5F). Serum ALT was also elevated in ob/ob mice, but it was unchanged after four weeks of siKHK treatment (Fig 5G). An improvement in metabolic dysfunction in ob/ob mice on fructose free diet prompted us to quantify the polyol pathway of endogenous fructose production. *Akr1b1* expression was unaffected in ob/ob mice and it did not change with T-KHK KD (Fig 5H). However, *Sord* was increased in ob/ob mice compared to controls, but it was not affected by T-KHK KD. In agreement with higher *Sord*, urinary fructose excretion was elevated in ob/ob mice (Fig 5I), consistent with increased endogenous fructose production and it was unaffected by T-KHK KD. Ob/ob mice were polyuric, due to glucosuria and produced dilute urine with low urine creatinine (Fig S7E). Urinary fructose corrected by urine creatinine was also elevated on ob/ob mice and it was unaffected by T-KHK KD (Fig S7F).

An increase in *Sord* and urinary fructose correlated with increased hepatic T-KHK (Fig 5J) and KHK-C (Fig 5K) in ob/ob mice on a fructose-free diet and a KD of T-KHK induced a complete loss of KHK. Next, we assessed DNL and ER stress pathways. Proteins mediating DNL, such as ACLY, ACC1, FASN and SCD1 were ~2 to 4-fold higher in ob/ob mice, compared to the controls (Figs 5J, S7G). Except for ACLY, KD of T-KHK had no effect on these DNL proteins. XBP1s was increased about 5-fold in ob/ob mice (Fig 5K), in line with increased ER stress in these mice (36). However, T-KHK KD did not alter XBP1s protein (Figs 5K, S7H). On the other hand, PERK Thr980 phosphorylation was markedly (p<0.0001) increase following T-KHK KD in ob/ob mice. Along with increased p-PERK, CHOP was 5-fold higher in ob/ob-siKHK, compared to the ob/ob-siCont group. In summary, KHK-C was increased in ob/ob mice likely due to increased endogenous fructose production. A KD of T-KHK in ob/ob mice resulted in metabolic improvements, which was associated with an improvement in p-PERK arm of ER stress, but not in DNL pathway.

### Knockdown of T-KHK in lipodystrophic mice improves metabolic dysfunction and ER stress, but not DNL

Following the exciting results in genetically obese ob/ob mice, we also confirmed increased KHK-C expression in livers of genetically obese leptin-resistant db/db mice (Fig S8A). Over time db/db, but not ob/ob, mice develop progressively more severe liver injury and we observed stepwise increased KHK-C in early, but not late stages of NAFLD. Rather than studying another model of genetic obesity, next we asked if T-KHK KD has a role in lean mice with metabolic dysfunction. We previously created a model of lipodystrophy by knocking out IR specifically in fat tissue (FIRKO). These mice completely lack visceral adipose tissue, but develop severe diabetes (37) and advanced NAFLD (38) since the liver becomes a primary site of fat storage. Two-month-old, chow-fed, FIRKO (IR flox/flox, cre+) and littermate control (IR flox/flox, cre-) mice were injected with T-KHK or luciferase control siRNA for four weeks. Lipodystrophic FIRKO mice gained similar amount of weight as littermate controls over four weeks, but a KD of T-KHK in FIRKO mice resulted in lower weight gain (Fig 6A). Most of the difference in weight can be attributed to a 25% reduced liver weight (4.0±0.2g) in FIRKO-siKHK (Fig 6B), compared to FIRKO-siCont mice (5.3±0.3g). Furthermore, FIRKO-siKHK group had lower hepatic TG and cholesterol, compared to FIRKO-siCont (Fig 6C). Random fed blood glucose was higher in FIRKO mice than in littermate controls and it markedly (p<0.001) improved following T-KHK KD (Fig 6D). Similarly, glucose tolerance was impaired in fasted FIRKO mice, which improved with T-KHK KD (Figs 6E, 6F). Serum TG (Fig 6G) and AST (Fig 6H) were elevated in FIRKO mice and decreased with T-KHK KD. Serum FFA and ALT were also elevated in FIRKO mice, but did not significantly improve with KD of T-KHK (Fig S8B). The polyol pathway in the liver was elevated in FIRKO mice, exemplified by increased *Akr1b1* and *Sord*, whereas T-KHK KD had no effect on these genes (Fig 6I).

**Figure 6.**
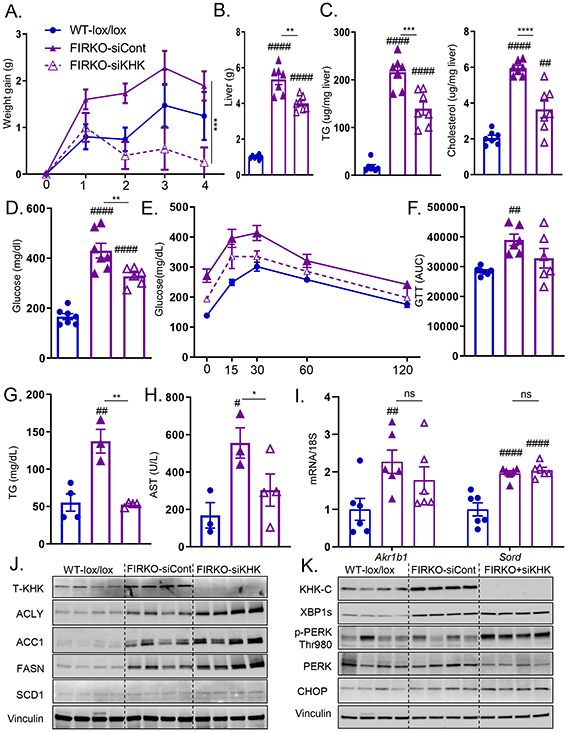
ER stress, but not lipogenesis, improves following T-KHK knockdown in lipodystrophic mice. Weight gain A) in 2-month-old, male, wild type, insulin receptor (IR) Lox/Lox, Adiponectin Cre negative (WT-lox/lox) littermates, as well as lipodystrophic, IR Lox/Lox, Adiponectin Cre positive, luciferase control injected (FIRKO-siCont) or KHK siRNA injected (FIRKO-siKHK) mice for four weeks, n=6-8 mice per group. B) Liver weight, hepatic C) triglycerides (TG) and cholesterol levels in these mice. Random fed D) blood glucose obtained at sacrifice, n=6-8 mice per group. E) Glucose tolerance test (GTT) in overnight fasted mice performed after 2 weeks following siRNA injection and F) area under curve (AUC) from GTT analysis. Serum G) TG and H) aspartate aminotransferase (AST) at sacrifice. Hepatic mRNA expression of I) genes regulating endogenous fructose production (*Akr1b1, Sord*). Western blot for KHK and proteins in the liver mediating J) *de novo* lipogenesis (ACLY, ACC1, FASN, SCD1) and K) ER stress (KHK-C, XBP1, PERK, CHOP) pathways. Sample number for WB is n=4 and for mRNA n=6 mice. Statistically significant results are marked with # using one-way ANOVA analysis, with Dunnett’s multiple comparisons test to the Chow group, # p<0.05; ## p<0.01; ### p<0.001; #### p<0.0001, while * represents a post-hoc analysis between the groups under the line, *p <0.05; ** p<0.01, *** p<0.001, **** p<0.0001. All data are presented as mean ± S.E.M.

Despite consuming a fructose-free diet, T-KHK (Fig 6J) and KHK-C (Fig 6K) were markedly elevated in FIRKO mice compared to the littermate controls, whereas T-KHK KD induced a complete loss of KHK. Proteins involved in DNL, such as ACLY, ACC1, FASN and SCD1 were ~2 to 5-fold higher in FIRKO mice, but KD of T-KHK actually increased these proteins in the liver (Figs 6J, S8C). XBP1s was ~3-fod higher in FIRKO mice compared to the control, but KD of T-KHK did not alter XBP1s (Figs 6K, S8D). On the other hand, PERK phosphorylation at Thr980 was 3-fold higher in FIRKO mice following T-KHK KD. In agreement with elevated p-PERK, CHOP was significantly (p<0.05) higher in FIRKO-siKHK, than in FIRKO-siCont group. In summary, KD of T-KHK in lipodystrophic FIRKO mice led to metabolic improvements, which was associated with an improvement in p-PERK arm of ER stress, but not in DNL pathway.

### KHK expression correlates with obesity and metabolic dysfunction in mice and early stages of NAFLD in humans

Due to an unexpected increase in KHK in the livers of mice with genetically induced obesity or metabolic dysfunction we asked if T-KHK expression correlates with these metabolic traits in Hybrid Mouse Diversity Panel (HMDP) (39). This study, conducted by Dr. Lusis group at the University of California, Los Angeles, more than 100 inbred strains of male or female mice were fed Western diet to induce various degrees of atherosclerosis and NAFLD (39). Hepatic microarray was performed and we correlated T-KHK expression with metabolic traits. T-KHK mRNA positively correlated with adiposity (R^2^= 0.292), serum insulin (R^2^=0.261), HOMA-IR (R^2^=0.267) and hepatic triglycerides (R^2^=0.176) in male mice (Fig 7A). Similarly, in female mice, T-KHK expression increased with increasing adiposity (R^2^=0.202), serum insulin (R^2^=0.233), HOMA-IR (R^2^=0.243) and hepatic triglycerides (R^2^=0.216) (Fig S9A). Next, we assess PERK arm of ER stress and found that *Eif2aka3* (R^2^=0.162), *Ddit3* (R^2^=0.305) as well as ER chaperons *Hspa5* (R^2^=0.198) and *Hsp90b1* (R^2^=0.231) correlated negatively with hepatic T-*Khk* expression in male mice (Fig S9B). This relationship was even stronger in female mice where *Eif2aka3* (R^2^=0.302), *Ddit3* (R^2^=0.349), *Hspa5* (R^2^=0.191) and *Hsp90b1* (R^2^=0.138) significantly (5.5e-22 for Ddit3) decreased with increasing T-*Khk* expression (Fig 7B). These results are in agreement with the data in Fig 1E showing that Ddit3, Hspa5 and Hsp90b1 were lower in HFD+F, compared to the Fruct group.

**Figure 7.**
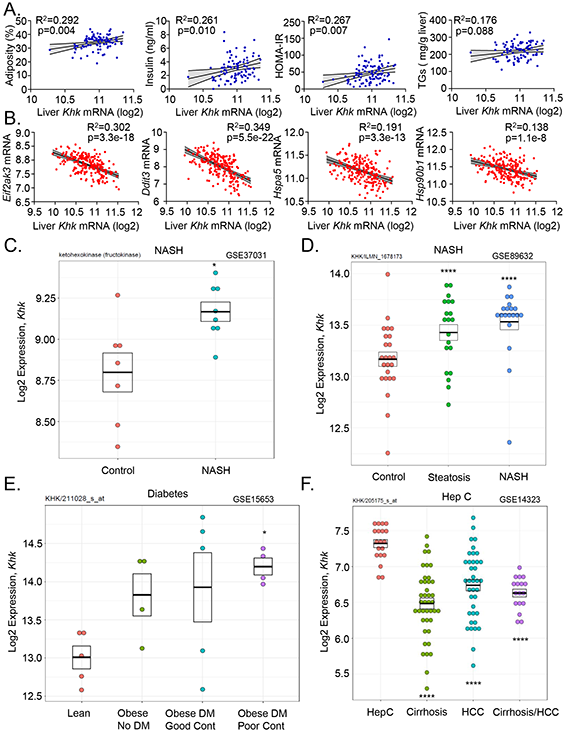
Hepatic ketohexokinase expression correlates positively with NAFLD and its risk-factors in mice and humans. The correlation between A) hepatic Khk expression and percent adiposity, serum insulin, HOMA-IR measure of insulin resistance and liver triglycerides (TG) in over 100 inbred strains of male (blue circles) male mice from Hybrid Mouse Diversity Panel. The correlation between B) hepatic Khk expression and PERK arm of ER stress (*Eif2ak3, Ddit3, Hspa5, Hsp90b1*). The correlation was determined by Pearson correlation coefficient using GraphPad Prism 8. Hepatic KHK expression in patients with C-D) NAFLD and NASH, as well as in patients with E) diabetes and F) hepatitis C infection. Human data were randomly searched and accessed from the studies that have deposited their data in a publicly available repository.

Since T-KHK expression correlated positively with NAFLD risk-factors in a large number of mice, next we asked if KHK expression correlates with NAFLD in human studies. Our bioinformatics group analyzed publicly available datasets from clinical studies involving patients with NAFLD. In a study by Lopez-Vicario et al. (40), T-*Khk* expression was increased in eight patients with morbid obesity and NASH compared to the controls (three cadaveric liver donors and four resection of metastasis to the liver) (Fig 7C). In a study by Arendt et al. (41), T-*Khk* mRNA was stepwise higher in 20 patients with steatosis and in 19 NASH patients, compared to 24 healthy controls (Fig 7D). In a study performed at our institution (Joslin Diabetes Center) (42), hepatic *Khk* expression increased in patients with obesity and type 2 diabetes, such that the obese patients with type 2 diabetes and poor glycemic control had the highest T-*Khk* expression (Fig 7E). On the other hand, T-*Khk* expression was not further elevated in patients with advanced and fibrotic NAFLD. In a study that had more rigorous phenotyping of NAFLD (43), where the patients were divided into four groups, T-*Khk* expression was stepwise higher starting with 14 healthy controls (NAS 0), 27 obese subjects (NAS 0), and 14 patients with steatosis (NAS 1-3), but not in 18 patients with advanced NASH (NAS 5-6) (Fig S9C). In agreement with this study, a study comparing 40 obese patients with mild NAFLD (fibrosis stage 0-1, 25% NAS≥5) to 32 BMI matched patients with advanced NAFLD (fibrosis stage 3-4, 56% NAS≥5), T-*Khk* expression actually decreased in the latter group (44) (Fig S9D). Consistent with decreased T-*Khk* expression in patients with advanced NAFLD, the patients with cirrhosis and/or hepatocellular carcinoma (HCC) secondary to chronic hepatitis C infection (45) had lower T-*Khk* expression than the patients with non-complicated chronic Hep C infection (Fig 7F).

In summary, *Khk* expression correlated positively with NAFLD risk-factors in over 100 inbred strains of male or female mice. Furthermore, in 241 patients with NAFLD and their controls, *Khk* expression was positively associated with obesity, metabolic dysfunction and early stages of NAFLD. However, KHK is not a marker of liver injury, as it decreased in patients with advanced NAFLD and in patients with severe liver disease secondary to hepatitis C infection.

## DISCUSSION

In the present study, we have explored the effects of a combined intake of sugar and a HFD on the development of severe NAFLD. We show that both nutrients can induce ER stress, whereas their combined intake results in unresolved ER stress accompanied by severe glucose intolerance, liver inflammation and hepatic fibrosis. Knockdown of T-KHK in mice consuming fructose on a HFD results in improved metabolic dysfunction via decreasing DNL and ER stress pathways. RNAseq analysis confirmed broad effects of T-KHK knockdown on ER stress and the hepatic transcriptome, far beyond what could be expected from its only known function to phosphorylate the simple sugar fructose. In line with its newly uncovered role, overexpression of KHK-C *in vitro* is sufficient to induce ER stress and impair autophagy. Increased KHK is observed in mice with genetically induced obesity or metabolic dysfunction consuming a fructose-free diet due to elevated endogenous fructose production. A knockdown of T-KHK in these mice improves metabolic dysfunction and restores the PERK arm of ER stress, without decreasing lipogenic enzymes. Consistent with its role in ER stress, hepatic *Khk* expression correlates positively with adiposity, serum insulin, HOMA-IR and hepatic triglycerides in over 100 inbred strains of male or female mice. Similarly, in 241 human subjects with NAFLD and their controls, *Khk* expression was upregulated in subjects with obesity, poor glucose control and early, but not late stages of NAFLD. While the main features of fructose are thought to be secondary to its lipogenic potential, here we describe a new role of fructose-induced KHK-C on ER stress that offers an explanation of how the combined intake of sugar and a HFD accelerates the development of NAFLD and metabolic complications.

Our work illustrates that consumption of both fructose and a HFD can upregulate XBP1s and PERK arms of ER stress. However, their combined intake leads to inadequate propagation of PERK signaling, resulting in unresolved ER stress. On the contrary, successful resolution of ER stress increases the expression of ribosomal subunits and expands ER protein folding capacity (46). Our RNAseq analysis revealed that a knockdown of T-KHK alters ER stress pathway. In cultured hepatocytes, ER stress can simply be triggered by overexpression of KHK-C. Indeed, our proteomic analysis revealed that ER mediated Regulation of Protein Complex Assembly is the most significantly upregulated pathway in KHK-C OE cells. While an increase in KHK-C is sufficient to upregulate ER stress, adding a second form of ER stress in terms of the combined HFD and fructose intake or serum starvation of KHK-C OE cells actually reduces BIP, a chaperone mediating resolution of ER stress. This suggest non-linear, dose-dependent nature of ER stress, where a single dose of ER stress triggers beneficial compensatory mechanism, whereas adding a second source of stress leads to unresolved ER strain and cellular injury. Indeed, unresolved ER stress is associated with a higher severity of NAFLD (47).

A novel role of KHK-C in mediating ER stress has not been previously described, but it is not completely unexpected. Fructose has been reported to induce ER stress (48) and conditions that induce *Khk-c* expression, namely hypoxia (12) and impaired glucose metabolism (13), are well-known triggers of ER stress. Here, we propose that dietary fructose serves as a physiologic stimulus to increase KHK-C and that elevated KHK-C then mediates ER stress leading to metabolic dysfunction attributed to dietary fructose. We show this to be the case for hepatic insulin resistance and steatosis. The combined intake of fructose and HFD leads to hepatic insulin resistance *in vivo* associated with a decrease in IRS-1 (49). A decrease in IRS-1 can be triggered *in vitro* by KHK-C induced perturbation in autophagy flux. Conversely, hyperinsulinemia equally downregulates IR in both HFD and HFD+F groups (50). Also, a decrease in ApoB observed in HFD+F group may contribute to severe hepatic steatosis in these mice. Indeed, ER mediated decrease in ApoB has been linked with liver lipid accumulation (33; 51). How KHK-C triggers ER stress is an area of future research. One hypothesis is that KHK-C mediates post-translational protein modifications leading to protein misfolding.

Our data suggest that signal transduction via the PERK arm of ER stress is sensitive to a degree of cellular stress. Impaired phosphorylation of translation initiation factor eIF2α is observed *in vivo* by the combined intake of fructose and HFD, as well as *in vitro* by OE of KHK-C and serum starvation. Phosphorylated eIF2 attenuates translation initiation and accelerates mRNA degradation (21). These effects are in agreement with our RNAseq analysis showing a profound impact of KHK KD on transcriptional regulation and nucleotide salvage pathways. Moreover, adequate eIF2α phosphorylation has been documented to protect hepatocytes from oxidative stresses induced by fructose metabolism (52). Mice with hepatocyte specific mutation in eIF2α, encoding phosphorylation S51A deficient protein, develop a more severe fatty liver disease when fed a high-fructose diet (52). Mechanistically, phosphorylated sugars have been proposed to occupy the catalytic site in the eIF2Bα subunit to modulate eIF2α phosphorylation (53). While eIF2α phosphorylation protects from nutrient-induced ER (54) and oxidative stress (55; 56), paradoxically, it can also induce cell death via strong and persistent induction of CHOP (57).

CHOP is universally recognized to mediate ER stress-induced apoptosis (58). What effect this plays in the setting of fatty liver disease is a matter of controversy. Some reports suggest that CHOP induction propagates liver inflammation, fibrosis, and oncogenesis (59; 60), while others document its protective role. First, CHOP is not increased in humans with NAFLD and NASH, in spite of increased eIF2α phosphorylation (61). Next, mice with CHOP deficiency develop more severe NAFLD with increased ER stress and hepatocyte apoptosis when fed a high fat, fructose and cholesterol diet (62). Additionally, an increase in CHOP is required to protect mice from another diet-induced liver injury, the methionine/choline deficient model (63). In our study, PERK/CHOP pathway is induced by HFD+F diet and by KHK-C OE, while a knockdown of KHK further upregulates PERK/CHOP pathway. In our study, an increase in CHOP following KHK KD is consistently associated with improved fatty liver disease. This is a more constant relationship than the improvement in lipogenesis induced by KHK KD. In fact, lipogenic enzymes are only decreased following T-KHK KD in mice consuming a large amount of fructose on a HFD. However, T-KHK KD does not reduce lipogenic enzymes in ob/ob or FIRKO mice on a fructose-free diet. This suggests that preventing fructose metabolism via a KD of T-KHK is required to downregulate DNL enzymes. On the other hand, KD of KHK improves metabolic derangements in ob/ob and FIRKO mice by mediating ER stress without decreasing DNL enzymes. Further, hepatic cholesterol is improved in all models following T-KHK KD, in line with the ER playing an integral part in cholesterol synthesis (64).

Consistent with unresolved ER stress, a combined intake of fructose and a HFD leads to the development of liver injury characterized by increased serum ALT, inflammation and hepatic fibrosis. A recent systemic review of 3,920 rodent models of NAFLD found that a high-fat, high-fructose diet indeed most closely resembles the severe liver injury observed in patients with NAFLD (19). Previously, we (23) and others (65–67) have showed that addition of sugar to a HFD propagates development of NAFLD more than the HFD alone. This may be secondary to increased lipogenesis, triggered by fructose metabolism and upregulation of KHK-C, which propagates ER stress-induced liver injury. Indeed, hepatic *Khk* expression correlates positively with markers of metabolic dysfunction in over 100 inbred strains of male or female mice. Moreover, hepatic *Khk* expression in these mice fed a Western diet is associated with decreased expression of PERK arm of ER stress and ER chaperones needed to resolve ER stress. Thus, KHK expression initially correlates with DNL and ER stress but with ongoing insult, ER stress response actually decreases. Non-linear, dose-dependent relationship between KHK and metabolic dysfunction is also observed in human livers. Khk expression is increased in early stages of NAFLD in 241 subjects with fatty liver disease and their controls. However, *Khk* expression did not correlate with more severe forms of liver injury. It appears that once liver injury sets in, ongoing upregulation of KHK is not necessary for continued liver damage. This is evident by persistently elevated XBP1s, which is not improved with a KD of T-KHK. Another possibility is that once patients develop severe liver disease their food intake decreases, which may translate into decreased KHK expression. However, in db/db mice hepatic KHK is initially elevated but then stabilizes, despite increasing severity of liver injury and ongoing obesity.

Here, we provide strong evidence that *Khk* expression correlates with multiple NAFLD risk-factors in mice and offer a mechanistic explanation underlying this interaction. Additionally, we provide, to date, the largest collection of data on *Khk* expression in human subjects with NAFLD. Prior studies have documented resolution of NALFD in mice with genetic knockout of KHK (66; 68; 69). This provides a strong rationale for targeting KHK for treatment of NAFLD. Several KHK inhibitors have already been developed (70–73), but this study points to additional questions that remain to be answered. Since KHK is upregulated in early stages of NAFLD, the efficacy of KHK inhibitors in models of advanced NAFLD needs to be determined. Long-term consequences of elevated CHOP in regards to liver carcinogenesis are unknown. Lastly, a profound effect of KHK KD on hepatic transcriptome calls for a more careful study of the potential side effects.

In conclusion, consumption of a simple sugar, fructose on a HFD accelerates development of metabolic complications and liver injury. While the role of fructose in supporting lipogenesis is well described, we report a novel function of fructose-induced upregulation of KHK-C in driving ER stress. This new role of KHK-C has a far-reaching impact as it profoundly affects global hepatic gene expression. Future studies are needed to broaden our understanding of this not so simple sugar metabolism.

## AUTHOR CONTRIBUTIONS

S.H.P., R.N.H, T.F, M.H.S, S.F. & S.S. conceptualized the study, performed the research, analyzed the data and wrote the manuscript; J.L.S.W, L.N, H.T & K.F performed the experiment in Fig 2. H.P & J.M.D analyzed RNAseq data and publically available biorepositories of human studies. J.B, J.R, C.D.K. & B.S. analyzed proteomics data in Fig 4. A.J.L & C.P provided and analyzed the data from MHDP study in Fig 7A, B. M.G & R.N.K provided data on db/db mice. P.A.K., S.D & C.R.K. helped guide the study, and assisted with preparation of the manuscript. All authors red and approved the manuscript before the submission.

## METHODS

### Animals and Diets

In accordance with NIH guidelines all animal protocols were approved by the IACUC of the University of Kentucky, Joslin Diabetes Center and Alnylam pharmaceuticals, where different experiments were performed. At Joslin Diabetes Center mice were housed at 20-22 ^0^C on a 12 h light/dark cycle with *ad libitum* access to food and water. C57BL/6J male mice at 10 weeks of age were purchased from Jackson Laboratory and fed either chow diet (Labdiet, Mouse Diet 9F) or high fat diet (Research diets, D12492) for 16 weeks. Caloric composition of the chow diet consisted of 23% protein, 21.6% fat and 55.4%carbohydrates, while the HFD had 20% protein, 60% fat and 20% carbohydrates. Mice were provided either tap water or 30% (weight/volume) fructose in water. The animals were weighed, and their food intake was recorded once per week. GTT and ITT were performed after 8 weeks on the diets. PTT was performed at 10 weeks on the diets. Mice were sacrificed from 8 to 11am, and one mouse from each cage, i.e. dietary group, was utilized before sacrificing the next mouse in the same cage. This was repeated until all four mice per cage were sacrificed. Male FIRKO mice were bred at Joslin Diabetes Center as previously described (38). At 2 months of age FIRKO mice (IR flox/flox, Adiponectin Cre +) were injected with KHK or luciferase (luciferase is not expressed in the mouse) control siRNA every two weeks for 4 weeks in total. Littermate IR flox/flox, Adiponectin Cre negative male mice were used as controls. Db/db mice were housed at Joslin Diabetes center and cared for by the lab members in Rohit Kulkarni lab. Db/db male mice were purchased from Jackson Laboratory at two months of age and they were fed chow diet for 4, 8 or 14 weeks to induce progressively more severe NAFLD. At Alnylam Pharmaceuticals, C57BL/6J male mice at 8-10 weeks of age were purchased from Charles River and fed either chow diet (Picolab® Rodent Diet 20 [5053] by LabDiet) or high fat diet (Research diets, D12492) supplemented with 30% fructose for 12 weeks. HFD+F mice were injected with KHK siRNA or PBS every two weeks for 8 weeks. The mice were sacrificed after 20 weeks on the diets. Ob/ob male mice were purchased from Jackson Laboratory at two months of age and provided regular chow diet (LabDiet 9F 5020, Cat# 0001329) at the University of Kentucky. This diet contains 0.98% sucrose. The mice were injected with KHK or luciferase control siRNA every two weeks for 4 weeks.

### Glucose, pyruvate and insulin tolerance test

For glucose tolerance test (GTT), mice were fasted overnight and injected intraperitoneally (IP) with 2g glucose per kg of body mass. Blood glucose levels were measured at 0, 15, 30, 60, and 120 minutes using a glucose meter (Infinity, US Diagnostics). Pyruvate tolerance test (GTT), was performed on overnight fasted mice and IP injected with 2g of pyruvate per kg of body mass. Insulin tolerance tests (ITT) were performed in nonfasted mice by i.p. injection of 1 mU insulin per kg of body mass. Blood glucose levels were measured at indicated times.

### Liver-specific KHK knockdown

Liver-specific knockdown was achieved by utilizing an siRNA conjugated to N-acetylgalactosamine (GalNAc). Alnylam Pharmaceuticals synthesized siRNA to specifically target mouse total Khk mRNA. The siRNA consists of 2 strands, guide and passenger. The guide strand carries the sequence information necessary for target-gene recognition, while the passenger strand supports loading into the RNA-induced silencing complex (RISC). siRNA has undergone chemical modifications to achieve long-lasting effect and specificity for hepatocytes. The combination of the 2′-fluoro, 2′-O-methyl and phosphorothioate modifications provides protection against exonuclease degradation, allowing for marked compound stabilization. The guide strand is conjugated to a trivalent GalNAc specifically recognized by the asialoglycoprotein receptor (ASGPR), which is highly expressed on the surface of hepatocytes, achieving hepatocyte-specific delivery and uptake. A GalNAc conjugate siRNA targeting luciferase was used as a negative control. Mice were injected subcutaneously with 20 mg/kg of KHK, 10 mg/kg luciferase control siRNA or PBS every two weeks depending on experimental design.

### Liver Histology, Sirius red and NAS scoring

Histology sections were prepared from formalin fixed paraffin embedded liver sections. H&E staining and Sirius red staining were performed by the University of Kentucky Pathology Research Core. Alnylam Pharmaceuticals Inc. completed in-house H&E staining on liver tissue from the experiment performed at their institution. Liver histology was graded by Experimental Pathology Laboratories, Inc. (EPL) to assign NALFD activity score. Our collaborators at Alnylam hired EPL to perform this work in an unbiased fashion on a fee-for-service basis.

### Assays quantifying serum and liver metabolic dysfunction

Serum samples were used for the quantification of triglyceride (Pointer scientific, T7532-1L) and cholesterol (Catachem, C104-01) following manufacturer’s guidelines. Plasma insulin level was quantified using ELISA kit (Crystal chem, 90080). ALT and AST level in the were measured using commercial ALT (Catachem, C164-0A), and AST (Catachem, C154-0A) kits.

Urinary fructose was quantified using a fructose assay kit (BioAssay Systems, EFRU-100). Liver lipid extracts were prepared as previously described (9). Liver homogenates were then used for the quantification of triglycerides and cholesterol described above. For the data in Figure 3 serum total, HDL and LDL cholesterol, AST, ALT, alkaline phosphatase and total bilirubin were measured at Alnylam Pharmaceuticals. In brief, Whole-venous blood was collected into serum separator tubes (BD Microtainer) and allowed to clot at room temperature for 30 min prior to centrifugation at 3000 RPM (1489 × g) for 10 min at 4 °C. Serum was then aliquoted and stored at −80 °C until analyses. Serum chemistries were analyzed using the AU400 chemistry analyzer (Beckman Coulter- Brea, CA, USA), with reagents provided by Beckman Coulter, Randox, and Sekisui Diagnostics. Differences between group means were evaluated for statistical significance using one-way ANOVA in GraphPad Prism 7 consistent with methodology previous published in (74).

### qPCR and mRNA quantification

mRNA was extracted by homogenizing liver tissue in trizol, treating with chloroform and participating in 70% ethanol. mRNA was purified using RNeasy Mini Kit columns (Qiagen, #74106). cDNA was made using High Capacity cDNA Revers Transcription Kit (Applied Biosystems, #4368813). qPCR was performed utilizing C1000 Thermal Cycler (BioRad, CFX384) and QuantStudio™ 7 Flex Real-Time PCR System (TermoFisher Scientific, 4485701). Mouse and human primer sequences used are listed in the supplemental tables 1 and 2, respectively.

### Protein Extraction and Immunoblot

Tissues were homogenized in RIPA buffer (EMD Millipore) with protease and phosphatase inhibitor cocktail (Bimake.com, B14002, B15002). Proteins were separated using SDS-PAGE and transferred to PVDF membrane (Millipore). Immunoblotting was achieved using the indicated antibodies listed in the supplemental table 3. Images were captured by ChemiDoc MP imaging system (BioRad, 12003154) and iBright imaging system (Thermo Fisher, CL1000). Quantification of immunoblots was performed using ImageJ.

### Generation of Stable KHK-C Overexpressing Hepatocytes

Human hepatic HepG2 cells were cultured under standard conditions in 1:1 Dulbecco’s Modified Eagle Medium (DMEM) and Ham’s F-12 media (Corning) supplemented with 10% fetal bovine serum (FBS, Corning) and 1% penicillin-streptomycin in a 5% CO2-humidified cell culture incubator at 37°C. Lentiviral particles were purchased from Origene (MR204149L2V) containing the mouse KHK-C sequence fused to GFP. HepG2 cells were transduced with lentivirus at a MOI of 5 and media was changed every two days thereafter (https://star-protocols.cell.com/protocols/913). After 5 days, cells were sorted for GFP-intensity and only individual GFP+ cells were collected in 96-well plates and propagated for an additional 2-3 weeks. Similarly GFP tagged KHK-C was overexpressed in mouse AML-12 cells. KHK-C was quantified in these clones by both QPCR and western blot. Only the highest expressing clones were used for subsequent experiments.

### Cell Culture Assays

For protein stability experiments, human wild type HepG2 and KHK-C OE cells were treated in low glucose media supplemented with 25 μg/mL cycloheximide, or media containing 25 mM fructose for 4- and 8 hours prior to protein extraction. For autophagy experiments, wild type and KHK-C OE HepG2 cells were cultured in complete media (DMEM/F12 with 10% FBS and 1% P/S), media without serum (DMEM/F12 with 1% P/S), or media supplemented with 50 μM chloroquine for 16 hours prior to protein extraction. Protein was used for subsequent western blot analysis.

### Proteomic analysis and bioinformatic data processing

Cell pellets of wild type and KHK-C OE AML-12 hepatocytes, with 4 replicates each, were solubilized in 700 μL of lysis buffer containing 8 M urea, 200 mM triethylammonium bicarbonate (TEAB), pH 8, 75 mM sodium chloride, 1 μM trichostatin A, 3 mM nicotinamide, and 1x protease/phosphatase inhibitor cocktail (Thermo Fisher Scientific, Waltham, MA), and sonicated using a probe sonicator. Protein concentrations were determined using a Bicinchoninic Acid Protein (BCA) Assay (Thermo Fisher Scientific, Waltham, MA). Proteins (2.1 mg) were reduced using 20 mM dithiothreitol (DTT) in 50 mM TEAB for 30 min at 37 °C, and after cooling to room temperature, alkylated with 40 mM iodoacetamide (IAA) for 30 min at room temperature in the dark. Samples were diluted 4-fold with 50 mM TEAB, pH 8, and proteins were digested overnight with a solution of sequencing-grade trypsin (Promega, San Luis Obispo, CA) in 50 mM TEAB at a 1:50 (wt:wt) enzyme:protein ratio at 37°C. This reaction was quenched with 1% formic acid (FA) and the sample was clarified by centrifugation at 2,000 x g for 10 min at room temperature. Clarified peptide samples were desalted with Oasis 10-mg Sorbent Cartridges (Waters, Milford, MA). 100 μg of each peptide elution were aliquoted, vacuum dried, and re-suspended in 0.2% FA in water at a final concentration of 1 μg/μL. Finally, indexed retention time standard peptides (iRT; Biognosys, Schlieren, Switzerland) (75) were spiked in the samples according to manufacturer’s instructions.

LC-MS/MS analyses were performed on a Dionex UltiMate 3000 system online coupled to an Orbitrap Eclipse Tribrid mass spectrometer (both from Thermo Fisher Scientific, San Jose, CA). The solvent system consisted of 2% ACN, 0.1% FA in water (solvent A) and 98% ACN, 0.1% FA in water (solvent B). Briefly, proteolytic peptides (400 ng) were loaded onto an Acclaim PepMap 100 C18 trap column (100 μm x 20 mm and 5 μm particle size; Thermo Fisher Scientific) for 5 min at 5 μL/min with 100% solvent A. Peptides were eluted on to an Acclaim PepMap 100 C18 analytical column (75 μm x 50 cm, 3 μm particle size; Thermo Fisher Scientific) at 0.3 μL/min using the following gradient of solvent B: 2% for 5 min, linear from 2% to 20% in 125 min, linear from 20% to 32% in 40 min, and up to 80% in 1 min, then holding at 80% for 10 min, and equilibrating the system at 2% B for 29 min – with a total gradient length of 210 min. All samples were acquired in data-independent acquisition (DIA) mode. Full MS spectra were collected at 120,000 resolution (AGC target: 3e6 ions, maximum injection time: 60 ms, 350-1,650 m/z), and MS2 spectra at 30,000 resolution (AGC target: 3e6 ions, maximum injection time: Auto, NCE: 27, fixed first mass 200 m/z). The DIA precursor ion isolation scheme consisted of 26 variable windows covering the 350-1,650 m/z mass range with an overlap of 1 m/z.

DIA data was processed in Spectronaut (version 15.1.210713.50606) using directDIA. Data was searched against a database containing all *Mus musculus* entries extracted from SwissProt (17,090 protein entries; January 27, 2022). Trypsin/P was set as digestion enzyme and two missed cleavages were allowed. Cysteine carbamidomethylation was set as fixed modification, and methionine oxidation and protein N-terminus acetylation as variable modifications. Data extraction parameters were set as dynamic. Identification was performed using 1% precursor and protein q-value. Quantification was based on the extracted ion chromatograms (XICs) of 3-6 MS2 fragment ions, local normalization was applied, and iRT profiling was selected. Differential expression analysis was performed using a paired t-test, and p-values were corrected for multiple testing, specifically applying group wise testing corrections using the Storey method (76). Protein groups with at least two unique peptides, q-value < 0.01 and absolute Log2(fold-change) > 0.58 were considered to be significantly altered.

Over-representation analyses (ORA) were performed using Consensus Path DB-mouse (Release MM11, 14.10.2021) (77) to evaluate which gene ontology terms were significantly enriched. Gene ontology terms identified from the ORA were subjected to the following filters: q-value < 0.05, term category = b (biological process), and term level > 3.

### Hybrid Mouse Diversity Panel

This study was performed by an independent investigator and they shared their banked data with us on 5/20/2021. In brief, over 100 inbred strains of male and female mice, from the Hybrid Mouse Diversity Panel study, were made hyperlipidemic by expressing the human ApoE-Leiden and CETP transgenes, and starting at 8-weeks of age were fed Western diet (Research Diets D10042101) for 16-weeks before sacrifice (39). Global transcript levels in aorta and liver were quantified, as well as complete metabolic profiling, including but not limited to adiposity, plasma lipid, insulin and glucose levels were measured. We correlated the expression of KHK in the liver with adiposity, insulin, HOMA-IR, and hepatic triglycerides (39). The correlation analysis was completed using the Pearson correlation coefficient.

### RNA Sequence Analysis and Bioinformatics Methods

HTG EdgeSeq mRNA sequence analysis was performed by the BioPolymers Facility at Harvard Medical School. Reads were aligned to the mouse transcriptome with Kallisto and the transcript counts were converted to gene counts with tximport. One sample was supposed to be KHK-treated, but its KHK expression was as large as the non-treated, and thus this sample was removed. Genes must have at least one count per million (CPM) in at least four samples, or they are filtered out. We normalized expression with trimmed mean of M-values (TMM) and transformed to logCPM with Voom. To discover the differential genes, we use limma, an R package that powers differential expression analyses. We applied the Fry function of the Rotation Gene Set Test (Roast) method in the limma R package to perform pathway analysis.

We obtained the Gene sets from the MSigDB Collections. We selected the gene sets that belong to the canonical pathways (CP) and Gene Ontology (GO). To get an overall view of the similarity and/or difference of the samples, we performed principal component analysis (PCA). In the volcano plot, the top genes with either the smallest p-values or largest logFC are labeled. In the heatmap, the same set of top genes are used, and the z-scores of the logCPM are plotted. We downloaded public datasets from the Gene Expression Omnibus (GEO) and compared between groups with limma.

### Statistical Analyses

All data are presented as mean ± SEM. The data analysis, comparing the effects of control to experimental conditions was first analyzed using one-way analysis of variance (ANOVA) with Dunnett’s multiple comparisons test for comparison of the individual groups. Significant difference among control and experimental groups is noted with a number sign (#) and significant difference between the individual groups under a black line is designated by an asterisk (*). For both signs, the single symbol represents a p value of <0.05, two symbols represent a p value of <0.01, and three symbols denote a p value of <0.001, and four symbols stand for a p value of <0.0001 throughout the study.

## DATA AVAILABILITY

Raw data and complete MS data sets were uploaded to the Center for Computational Mass Spectrometry, the MassIVE repository at UCSD, and can be downloaded using the following link: ://massive.ucsd.edu/ProteoSAFe/dataset.jsp?task=832a44837fba49219ebef460ef6de77c MassIVE ID number: MSV000089043; ProteomeXchange ID: PXD032190.

(Go to *https://massive.ucsd.edu/ProteoSAFe/private-dataset.jsp?task=832a44837fba49219ebef460ef6de77c* in the upper right corner: Enter Username: MSV000089043_reviewer; Password: winter In the repository entry in MassIVE click the ‘Browse Dataset Files’ button.)

RNAseq data will be deposited in a public repository after acceptance of the manuscript for publication.

