## Supplemental Figures for "Fructose Induced KHK-C Increases ER Stress and Modulates Hepatic Transcriptome to Drive Liver Disease in Diet-Induced and Genetic Models of NAFLD"

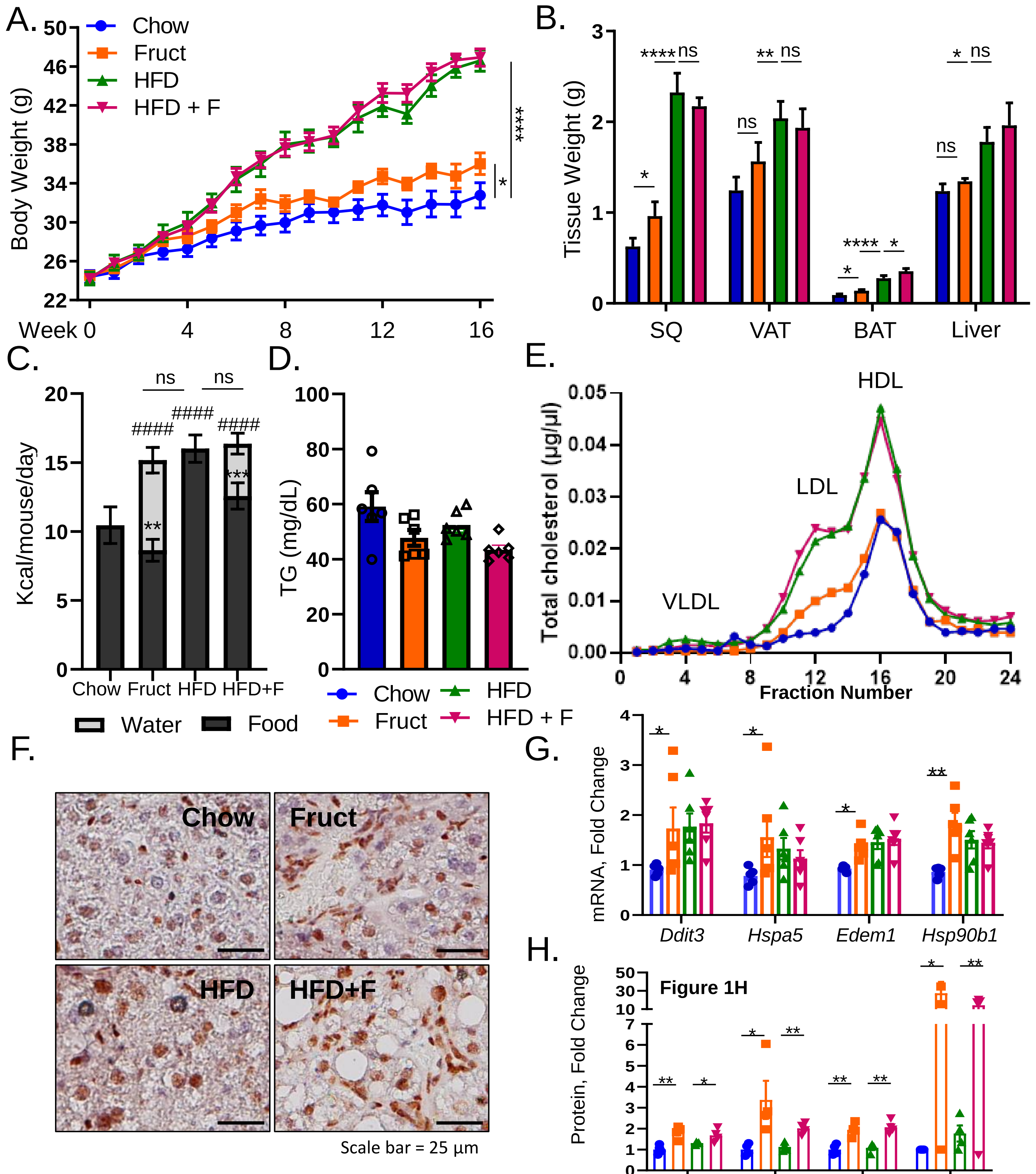


### Supplemental Figure 1

Weight gain A) in male, C57BL/6J, mice drinking regular water (Chow) or 30% fructose-sweetened (Fruct) water on Chow diet, as well as high fat diet fed mice on regular (HFD) or fructose water (HFD+F) for 16 weeks. n=6-8 mice per group. Weights of subcutaneous (SQ), visceral adipose tissue (VAT), brown adipose tissue (BAT) and liver at sacrifice. C) Food and water intake were measured once per week for the duration of the experiment. D) Serum triglycerides (TGs) at sacrifice. E) Total cholesterol measured following FPLC fractionation of serum. F) Immunohistochemistry for pPERK in livers of these mice. G) mRNA expression of ER stress markers. H) Image J quantification of WB from Figure 1G. All data present mean ± S.E.M.

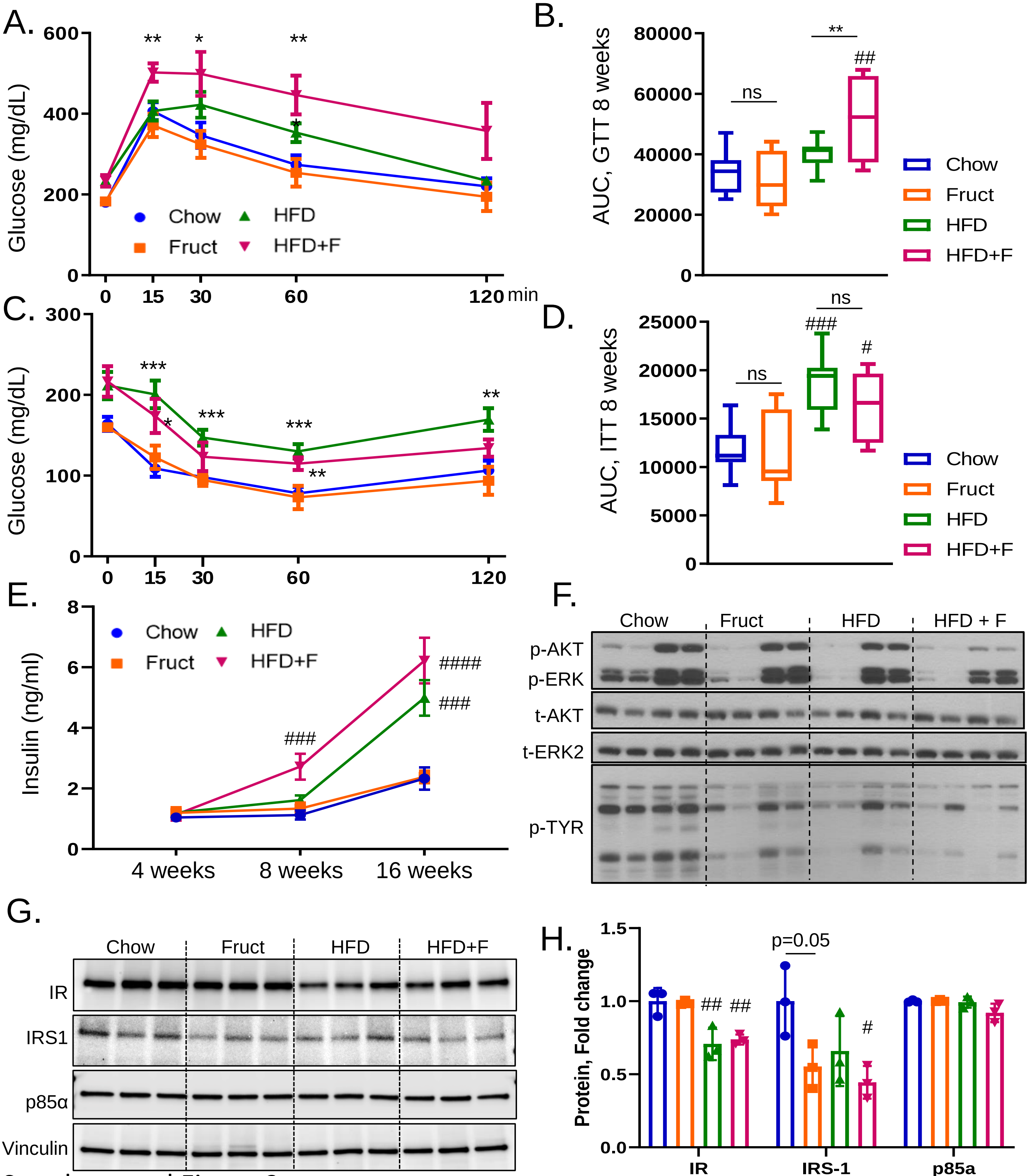


#### Supplemental Figure 2

A) Glucose tolerance test (GTT) following 2h fast after 8 weeks on the diets. n=6-8 mice per group. B) Area under curve (AUC) quantification of GTT data in A. C) Insulin tolerance test (ITT) performed after 9 weeks on the diets. D) AUC quantification of ITT data in C. E) Overnight fasted insulin levels measured after 4, 8 and 16 weeks on the diets. Blood was collected via tail vein. Western blot of F) phospho and total Akt, ERK and pan tyrosine phosphorylation from liver homogenates. G) Western blot of insulin signaling molecules (IR, IRS1, p85α) from liver homogenates. H) Image J quantification of WB data in G. Sample number for WB is n=4 . All data are presented as mean ± S.E.M.

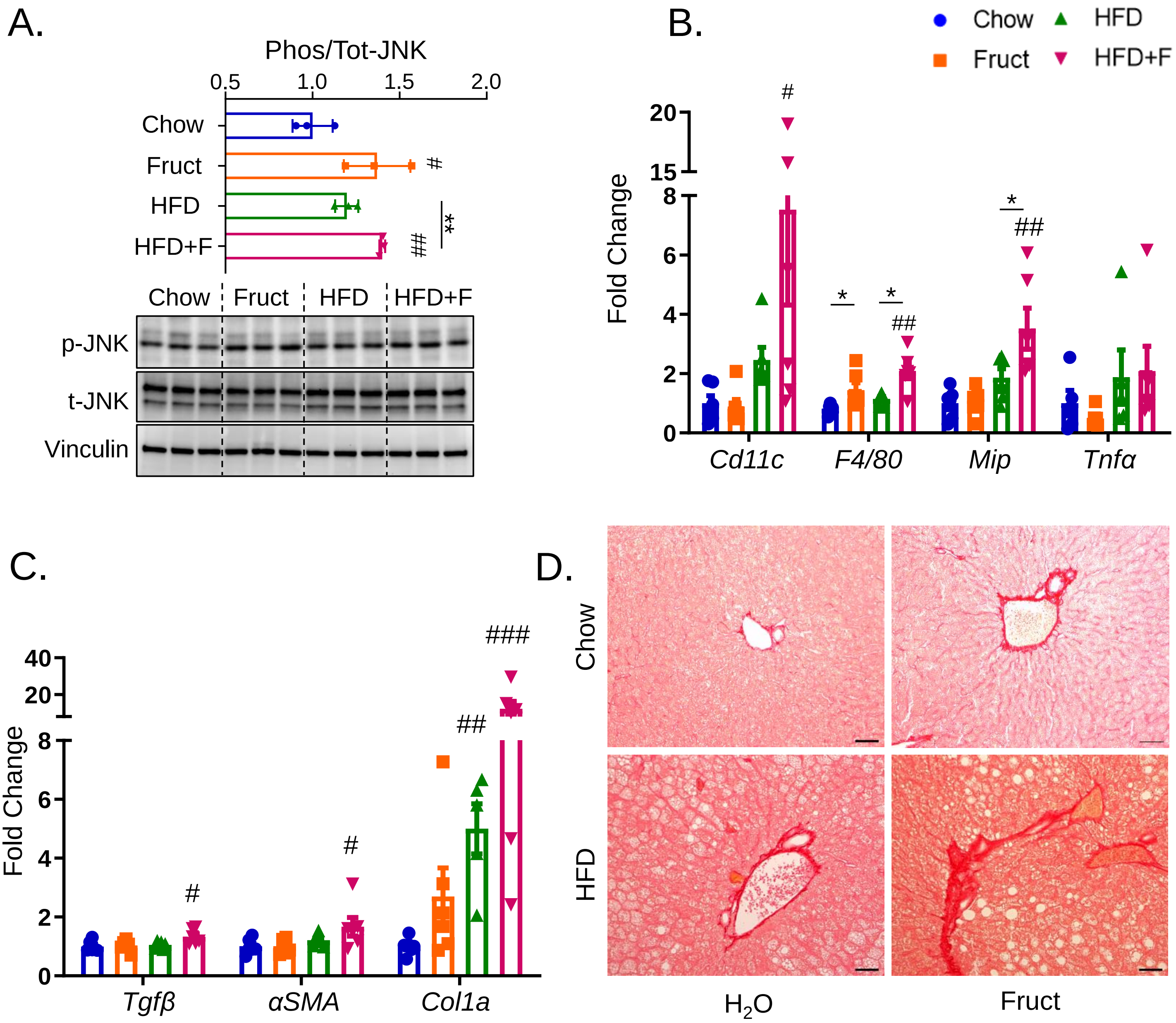


#### Supplemental Figure 3

A) Western blot (WB) analysis of phosphorylated and total c-Jun NH2-terminal kinase (JNK) from the livers of male, C57Bl6/J mice drinking regular water (Chow) or 30% fructose-sweetened (Fruct) water on Chow diet or regular (HFD) or fructose water (HFD+F) on a HFD for 16 weeks. WB quantification was preformed using image J software. n=3 mice per group. Hepatic mRNA expression from these mice of genes mediating B) inflammation (*Cd11c, F4/80, Mip,Tnfα,*) and C) collagen deposition (*Tgfβ*, *αSMA*, *Col1a*). n=6 mice per group. All data are presented as mean ± S.E.M. D) Representative images of Sirius Red stained liver sections from these mice. Scale bar represents 20 μm.

### Supplemental Figure 4

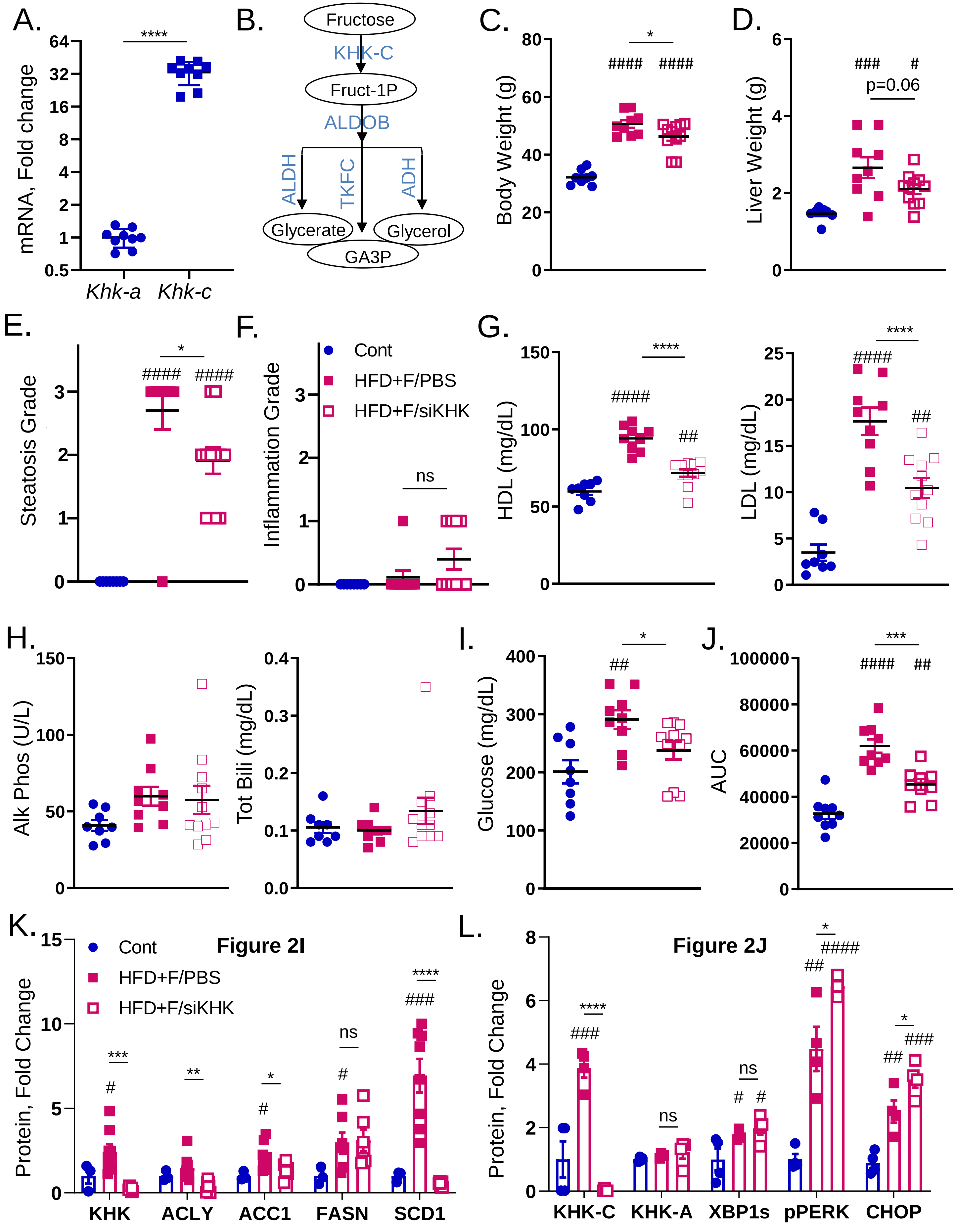

### Supplemental Figure 4

A) Hepatic mRNA expression of *Khk-a* and *Khk-c* isoforms in male, C57BL/6J mice drinking regular water on Chow diet. B) An illustration of fructolysis pathway representing the first, second and third steps of fructose metabolism. C) Body and D) liver weight of Chow or HFD-fed mice supplemented with 30% fructose for 20 weeks and injected with PBS (HFD+F/PBS) or total KHK siRNA (HFD+F/siKHK) for the last 8 weeks of the experiment. E) Steatosis score and F) Inflammation score based on liver histology as graded by a blinded expert pathologist from Experimental Pathology Laboratories, Inc. Serum G) High-density lipoprotein (HDL), and low-density lipoprotein (LDL), H) alkaline phosphatase (Alk Phos) and total bilirubin (Tot Bili) from these mice at the time of the sacrifice. I) Basal glucose levels and J) area under curve (AUC) calculated from glucose tolerance test (Fig 2H) from these mice obtained after 18 weeks on the diets. n= 8 to 11 mice per group. Image J quantification of western blot data presented in K) Figure 2I and L) Figure 2J. n= 3 to 8 mice per group. All data are presented as mean ± S.E.M.

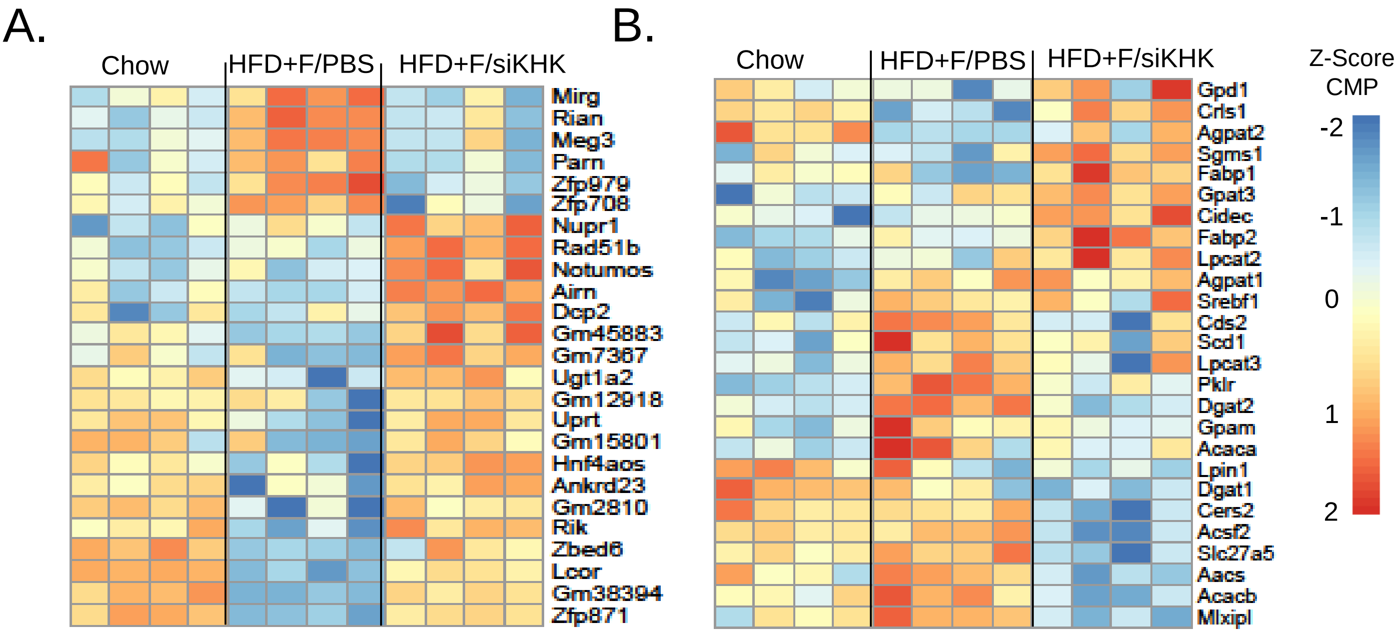


### Supplemental Figure 5

Heatmap representation of RNAseq data showing A) the most altered genes in HFD+F-fed mice treated with PBS or KHK siRNA and B) the expression of genes involved in de novo lipogenesis pathway. Chow control and HFD fed mice supplemented with 30% fructose for 20 weeks were injected with PBS (HFD+F/PBS) or total KHK siRNA (HFD+F/siKHK) for the last 8 weeks on diets. The scale represents clipped Z-scores of counts per million (CMP) RNAseq reads. n=4 mice per group.

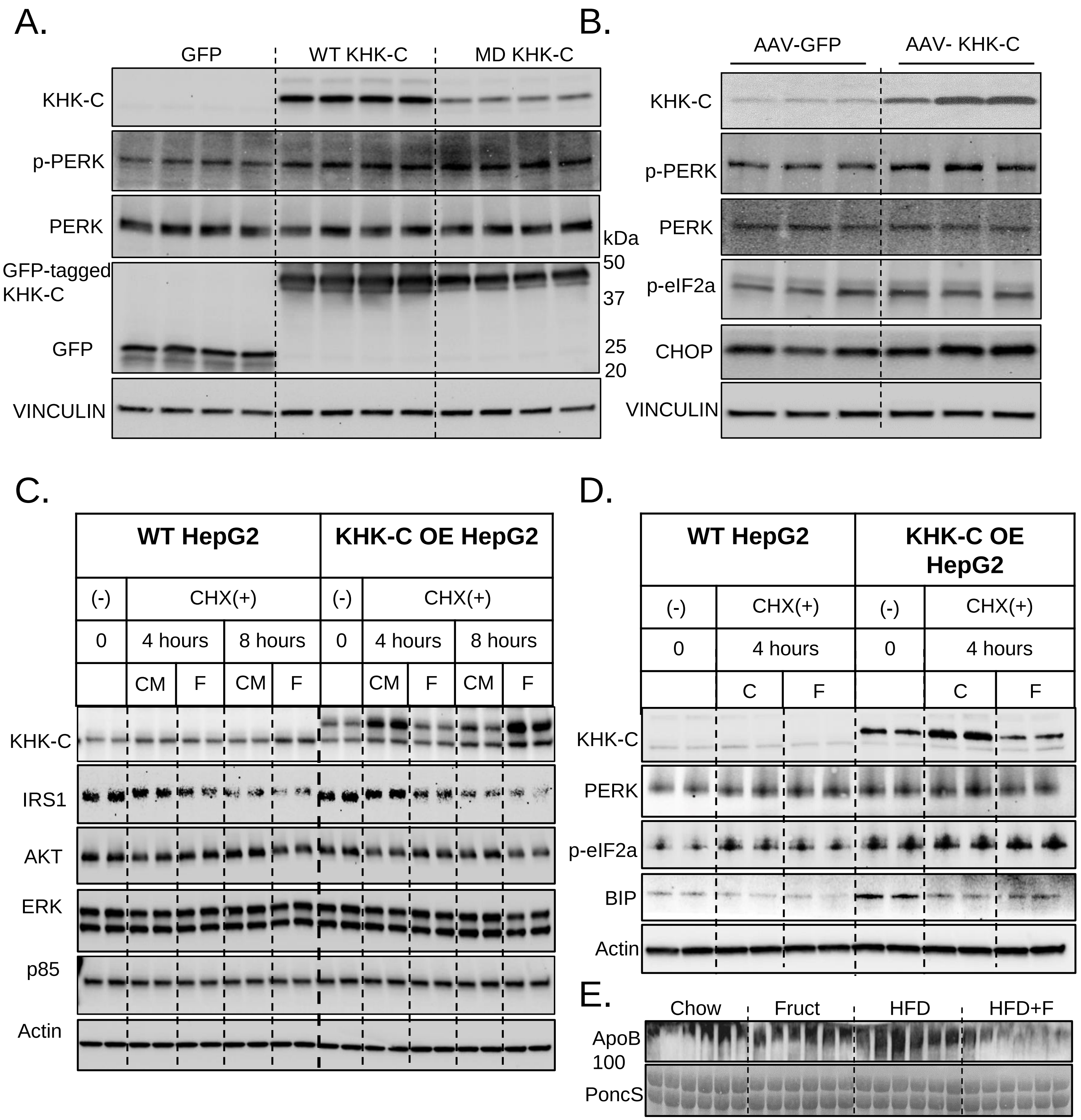


#### Supplemental Figure 6

HepG2 cells were treated with lentivurs caring either GFP, GFP tagged wild type KHK-C (WT KHK-C) or GFP tagged with mutant kinase dead KHK-C (MD KHK-C). GFP positive cells were sorted and cultured in fructose-free, DMEM media supplemented with 10% FBS. A) WB from these cells. n=4 wells per group. B) Male, C57BL/6J mice were injected with AAV-8 carrying GFP (AAV-GFP) or wild type KHK-C (AAV-KHK-C) under TBG promoter. The virus was injected via tail vain and the mice were on chow diet for one week. Livers were harvested for WB analysis. n=3 mice per group. C) Standard HepG2 cells (WT HepG2) and wild type GFP tagged KHK-C OE HepG2 cells (KHK-C OE HepG2) were treated with culture media (CM) or a combination of CM and 25 mM fructose (F) with ± cycloheximide (CHX) for 4 and 8 hours. Western blot was used to analyze C) insulin signaling and D) ER stress pathways. n=2 mice per group. E) Western blot analysis of serum samples for ApoB 100 using mice from the Fig 1. PonceauS (PoncS) was used as a loading control. n=6 serum samples per group.

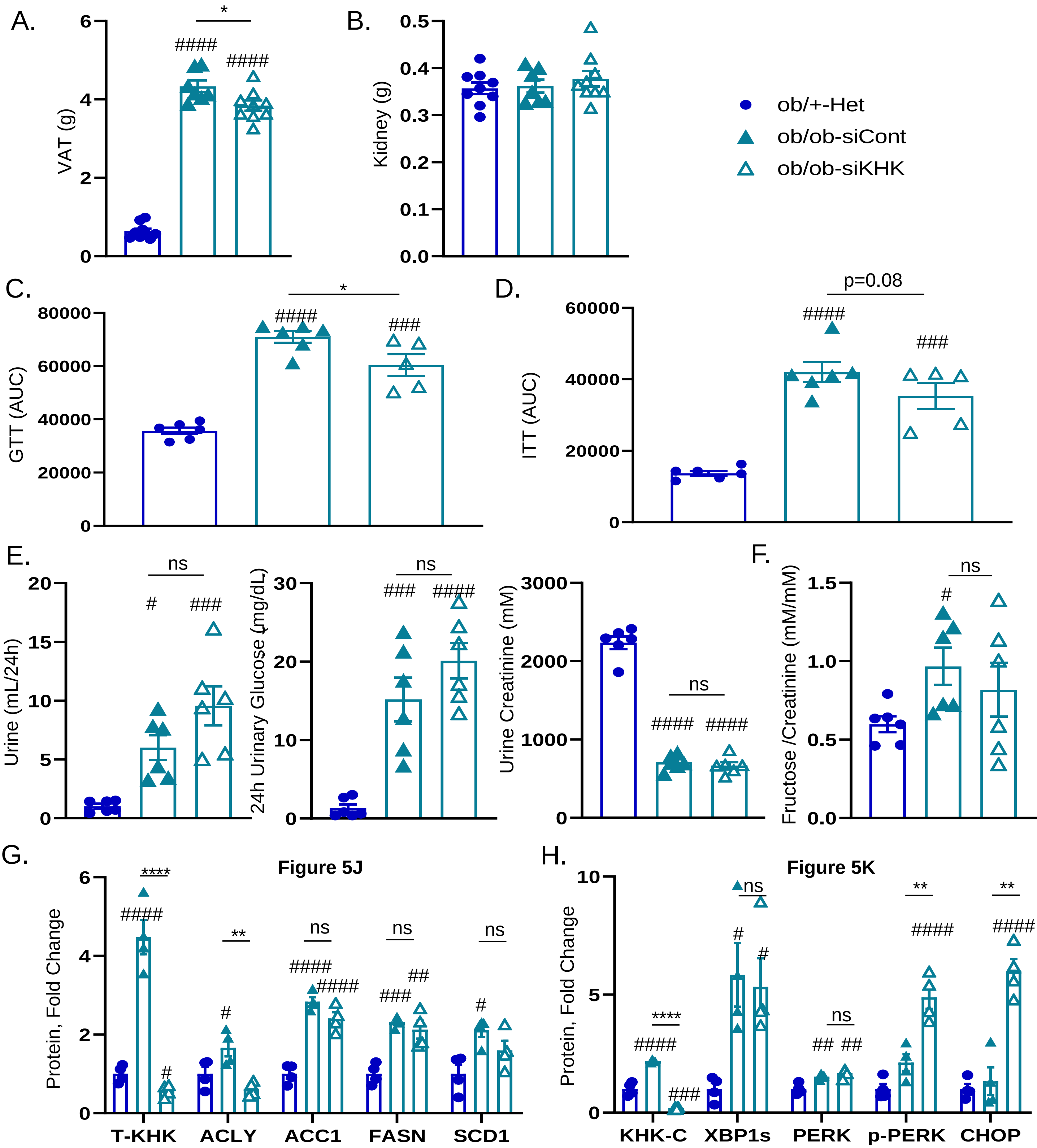


### Supplemental Figure 7

A) Visceral adipose tissue (VAT) and B) Kidney weight from 2-month-old, male, heterozygote (Ob/+-Het), as well as in homozygote, leptin-deficient mice (Ob/Ob-siCont) treated with luciferase control, or KHK siRNA (Ob/Ob-siKHK) for four weeks, n=6-9 mice per group. C) Area under cure (AUC) calculated from glucose tolerance test (Fig 5D) performed after 2 weeks following siRNA injection. D) AUC calculated form insulin tolerance test (Fig 5E) performed after 3 weeks following siRNA injection. E) Urine volume collected over 24h. Urinary glucose and Creatinine concentrations determined from 24h urine collection. F) Urinary fructose to creatinine ratio. Image J quantification of western blot data G) presented in Figure 5J and H) Figure 5K. n= 4 mice per group. All data are presented as mean ± S.E.M.

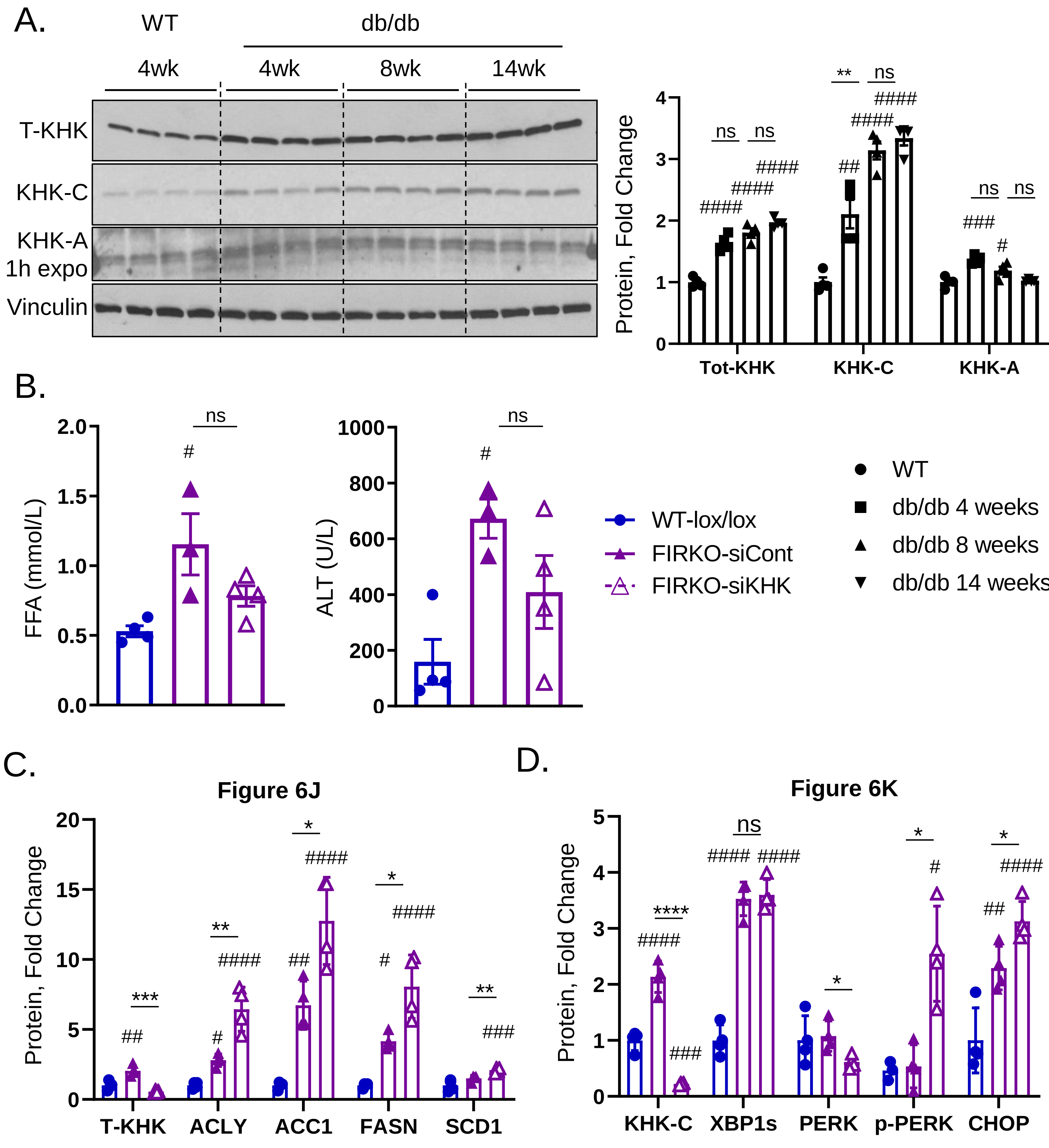


#### Supplemental Figure 8

A) Western blot (WB) analysis of hepatic total KHK (T-KHK) as well as KHK-A and KHK-C isoforms from leptin receptor deficient (db/db) and db/+ heterozygote control (WT) mice after 4, 8 and 14 weeks on chow diet. Image J quantification of WB data. n= 4 mice per group. B) Serum free fatty acids (FFA) and alanine aminotransferase (ALT) from 2-month-old, male, wild type, insulin receptor (IR) Lox/Lox, Adiponectin Cre negative (WT-lox/lox) littermates, as well as lipodystrophic, IR Lox/Lox, Adiponectin Cre positive, control mice injected with luciferase control (FIRKO-siCont) or KHK siRNA (FIRKO-siKHK) for four weeks, n=6-8 mice per group. Image J quantification of western blot data C) presented in Figure 6J and D) Figure 6K. n= 4 mice per group. All data are presented as mean ± S.E.M.

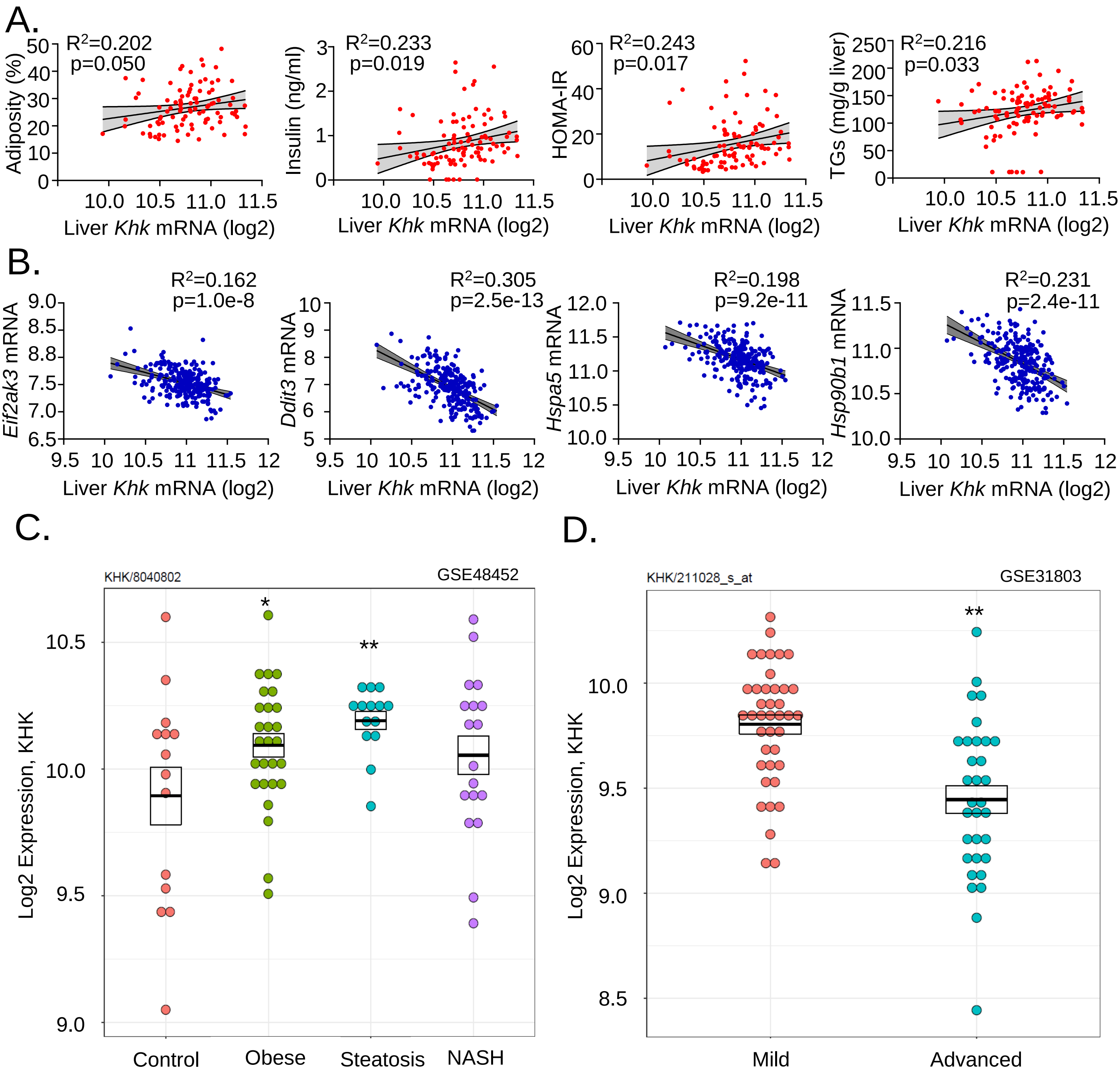


### Supplemental Figure 9

A) Hepatic total Khk expression correlated positively with adiposity, serum insulin, HOMA-IR and liver triglycerides (TG) in strains of female (red dots) mice from Hybrid Mouse Diversity Panel study. B) Hepatic total Khk expression correlated negatively with PERK arm of ER stress in strains of male (blue dots) mice from the same study. C) Hepatic mRNA expression of total Khk from 14 healthy controls (NAFLD Activity Score=NAS 0), 27 obese subjects (NAS 0), 14 patients with steatosis (NAS 1-3), and in 18 patients with advanced NASH (NAS 5-6). D) Hepatic mRNA expression of total Khk from 40 obese patients with mild NAFLD (fibrosis stage 0-1, 25% NAS≥5) and 32 BMI matched patients with advanced NAFLD (fibrosis stage 3-4, 56% NAS≥5). Human data were searched and accessed from the studies that have deposited their data in a publicly available repository. The data were analyzed by bioinformatics core at Joslin Diabetes Center.
